# Roles of adenine methylation and genetic mutations in adaptation to different temperatures in *Serratia marcescens*

**DOI:** 10.1101/822080

**Authors:** Matthieu Bruneaux, Ilkka Kronholm, Roghaieh Ashrafi, Tarmo Ketola

**Affiliations:** Department of Biological and Environmental Science, University of Jyväskylä, Finland

**Keywords:** Experimental evolution, Methyltransferase, single molecule real-time sequencing

## Abstract

Epigenetic modifications can contribute to adaptation, but the relative contributions of genetic and epigenetic variation are unknown. Previous studies on the role of epigenetic changes in adaptation in eukaryotes have nearly exclusively focused on cytosine methylation (m5C), while prokaryotes exhibit a richer system of methyltransferases targetting adenines (m6A) or cytosines (m4C, m5C). DNA methylation in prokaryotes has many roles, but its potential role in adaptation still needs further investigation. We collected phenotypic, genetic, and epigenetic data using single molecule real-time sequencing of clones of the bacterium *Serratia marcescens* that had undergone experimental evolution in contrasting temperatures to investigate the relationship between environment and genetic, epigenetic, and phenotypic changes. This data provided a detailed description of the methylation landscape of *S. marcescens* and allowed us to examine the potential contributions of genetic and epigenetic changes to phenotypic adaptation. The genomic distribution of GATC motifs, which are the main target for m6A methylation, and of partially methylated epiloci pointed to a link between m6A methylation and regulation of gene expression in *S. marcescens*. Evolved strains, while genetically homogeneous, exhibited many polymorphic m6A epiloci. There was no strong support for a genetic control of methylation changes in our experiment, and no clear evidence of parallel environmentally-induced or environmentally-selected methylation changes at specific epiloci was found. Both some genetic and epigenetic variants were associated with some phenotypic traits. Overall, our results suggest that both genetic and adenine methylation changes have potential to contribute to phenotypic adaptation in *S. marcescens*, but that any environmentally-induced epigenetic change occurring in our experiment would probably have been quite labile.

## Introduction

While the traditional view of evolution is that adaptation proceeds via changes in DNA sequences, any heritable phenotypic variation can in principle be subject to natural selection. In recent years, some epigenetic changes such as DNA methylation changes have been found to be heritable and could thus theoretically play a role in evolution (Jablonka and Raz, 2009; Day and Bonduriansky, 2011; Danchin et al., 2011; Kronholm and Collins, 2016; Danchin et al., 2019). However, even if convincing cases of epigenetic inheritance have been documented (Cortijo et al., 2014), the role of epigenetic variation in evolution has been met with some skepticism (Charlesworth et al., 2017), the main argument for caution being that despite frequent observations we know very little about the relative contributions of genetic and epigenetic variation to adaptation. Addressing this question requires the simultaneous characterization of genetic, epigenetic and phenotypic changes in a population, which remains extremely challenging in a natural system, and so far empirical results have lagged behind theoretical models. Experimental evolution under controlled laboratory conditions with bacteria that have short generation times and large population sizes provides the opportunity to study this question in a practical way.

Non-genetic inheritance encompasses various mechanisms, such as structural inheritance via propagating templates (e.g. prions), self-sustaining regulation loops and DNA methylation (Jablonka and Raz, 2009). DNA methylation is particularly well studied and exists both in eukaryotes and in prokaryotes. While DNA methylation can have phenotypic effects in both of those taxonomic groups, important differences exist between them. In eukaryotes, C5-methylcytosine (m5C) is the best known modification and occurs in CpG-rich regions. Genome methylation patterns are maintained after DNA replication via maintenance methyltransferases but DNA methylation can be reset in the germ line. In prokaryotes, N4-methylcytosine (m4C) and N6-methyladenine (m6A) are known in addition to m5C (Ratel et al., 2006; Casadesús and Low, 2006). Methylation is lost on the newly synthetized DNA strand during replication, and methylation restoration depends on the activity of sequence-specific methyltransferases after passage of the replication fork.

DNA methylation systems in prokaryotes are very diverse and include restriction-modification (RM) systems and orphan methyltransferases (MTases). RM systems associate an MTase activity and a complementary endonuclease activity: methylation of the target sequence protects DNA from restriction. Orphan MTases do not have an associated restriction enzyme but are thought to be evolutionary derived from RM systems. DNA methylation in bacteria has a wide range of well-documented roles: protection against foreign DNA via RM systems, regulation of DNA replication and mismatch repair based on hemimethylated DNA, control of mobile elements and regulation of gene expression (Casadesús and Low, 2006; Sánchez-Romero et al., 2015). Crucially, DNA methylation patterns can be inherited in prokaryotes: orphan MTases can compete with other regulatory proteins for access to cognate DNA sequences, for example in promoter and regulatory regions, resulting in self-regulating loops that can be maintained across cell division (Braaten et al., 1994).

We used *Serratia marcescens* as a bacterial model in evolution experiments and characterized genetic, epigenetic and phenotypic differences between strains evolved from a common ancestor culture. *S. marcescens* is an opportunistic pathogen which can live in a variety of environments, including inside insect and mammalian hosts, with an optimal growth temperature close to 31 *^◦^*C (Grimont and Grimont, 1978; Flyg et al., 1980; Ketola et al., 2013). *S. marcescens* possesses a Dam MTase targeting adenines in 5*^’^*-GATC-3*^’^* motifs, and it has been shown that this enzyme is important to enable the mismatch repair machinery to distinguish between the original and the new DNA strand during replication (Ostendorf et al., 1999; Blow et al., 2016).

While examples of inherited epigenetic changes and their involvement in bacterial adaptation exist (Adam et al., 2008; Atack et al., 2015), we do not fully understand yet the relative importance of epigenetic and genetic variation in evolution. Here, we studied the role of epigenetics in evolution in a prokaryotic system by (1) characterizing the epigenetic variation related to m6A in *Serratia marcescens* after evolution in different conditions, (2) determining if the observed epigenetic variation was associated with specific environmental conditions, and (3) comparing the potential contributions of epigenetic and genetic variation to phenotypic variation.

We addressed these questions by phenotyping and single-molecule real-time (SMRT) sequencing of bacterial strains from an evolution experiment conducted at three contrasting thermal regimes: 31 *^◦^*C constant, 38 *^◦^*C constant, and 24–38*^◦^*C fluctuating environments (Ketola et al., 2013). By introducing a common garden step before phenotypic measurements and DNA sequencing, we focused on the identification of evolutionary differences in methylation patterns which would be stable enough to be propagated across several generations after their establishment once selection pressure was relieved and we ignored epigenetic mutations with fast back-mutation rates which would be more akin to phenotypic plasticity. Additionally, to investigate the mechanism by which adenine methylation could affect cell processes in *S. marcescens*, we examined if the genomic distribution of the GATC cognate motif exhibited any informative pattern.

## Materials and Methods

### Experimental evolution and phenotypic measurements

We used bacterial strains that were obtained from a previous evolution experiment (Ketola et al., 2013). To summarize this experiment briefly, we let populations of *Serratia marcescens* initiated from a single common ancestor colony evolve under either constant or fluctuating temperatures during three weeks (treatments: constant 31 *^◦^*C, constant 38 *^◦^*C or alternating daily between 24 *^◦^*C and 38 *^◦^*C) (Ketola et al., 2013) (Supplementary Figure S1). Note that lines evolved in 38 *^◦^*C were not reported in Ketola et al. (2013). The experiment lasted approximately 70 generations (based on 1*/*10 daily subcultures for 3 weeks). Given that population sizes were of the order of 8 *×* 10^6^ cells in each 400 ➭L culture well at plateau, and that the fraction transferred to the next generation was 1*/*10 of the previous culture, the effective population size is estimated to be *N_e_* = 2.6 *×* 10^6^ during the experiment (using *N_e_* = *N*_0_*g* from Lenski et al. (1991), where *N*_0_ = 8 *×* 10^5^ cells is the bottleneck size and *g* = 3.3 is the number of generations between transfers). After experimental evolution, individual colonies were isolated from each population by plating and stored at *−*80 *^◦^*C in 50 % glycerol.

We randomly selected one frozen clone from each population for sequencing (10 from the 31 *^◦^*C treatment, 8 from the 38 *^◦^*C treatment and 10 from the fluctuating treatment, i.e. 28 evolved strains sequenced in total). As a reference and since the frozen stocks for the common ancestor itself could not be revived successfully prior to sequencing, we also sequenced the stock strain from ATCC (ATCC 13880) from which the common ancestor colony was derived (Supplementary Figure S1). This strain is referred to as the “reference strain” in the rest of this manuscript. We used the phenotypic data from Ketola et al. (2013) (Supplementary Figure S2), which included measurements of growth rate and yield at constant 24 *^◦^*C, 31 *^◦^*C and 38 *^◦^*C for the evolved strains. In addition, growth rate and yield had been measured in a series of novel environments: under redox balance stress (1 mg*/*ml dithiotreitol), in the presence of the ciliate predator *Tetrahymena thermophila* and in the presence of a virus, the lytic bacteriophage PPV. For further details on these phenotypic measurements, see Ketola et al. (2013).

### Sequencing and genome annotation

#### Sequencing

We used single molecule real-time sequencing (SMRT) with the PacBio platform to sequence the evolved strains and the reference strain. Since no template amplification takes place prior to sequencing on a PacBio platform in order to detect base modifications, relatively large amounts of DNA per strain are necessary with this method. Evolved strains were thawed and grown overnight in 150 ml of liquid medium and DNA was extracted using the Wizard Genomic DNA Purification Kit from Promega (WI, USA). One DNA sample (20 to 60 µg) per strain was sequenced by the DNA Sequencing and Genomics Laboratory of the University of Helsinki on a PacBio RS II sequencing platform using P6-C4 chemistry. Two single-molecule real-time sequencing (SMRT) cells were run per DNA sample.

PacBio software and recommended protocols were used with default parameters for the assembly and modification calling pipeline. For each strain, reads from the RS II instrument were assembled with PacBio `RS_HGAP_Assembly.3`, as implemented in SMRTportal 2.3.0. The resulting assembly for each strain was processed with the Gap4 program to generate *de novo* a first draft sequence for this strain and to circularize it. PacBio `RS_Resequencing.1` protocol was then run 2 to 3 times for each sample to map the reads to the draft sequence and generate a consensus sequence for each strain. The average coverage of the draft chromosome per strain was high (from 102 to 413, average 268 across all strains). We used Samtools (version 1.7, Li et al. (2009)) to examine PacBio reads mapped back to their respective assemblies in order to check for evidence of polymorphism within each sequenced strain. Such polymorphism would indicate that a given strain would be comprised of several genetic clones, which is possible if an isolated colony grew from more than a single isolated cell. Based on the Samtools pileup files, the distribution of proportions of mismatches for each position within a strain was compatible with the expected base error rate for PacBio SMRT sequencing (about 11-15%, Rhoads and Au (2015)), except for a short ca. 700bp region comprised of repeated sequences and which was ignored in downstream analyses. Given the high coverage of the chromosome sequences across all strains, the base calling was considered accurate outside of this repeat region and genetic variants were directly called from an alignment of the 29 assembled chromosomes built with Mugsy (Angiuoli and Salzberg, 2011).

### Estimation of methylated fractions

Inter-pulse duration (IPD) ratios were used with the PacBio `RS_Modification` and `Motif_Analysis` protocol to detect modified bases and identify methylation sequence motifs. Methylated adenines (m6A) are the modified bases with the strongest kinetic signal (Clark et al., 2012) and we only analysed those modifications in downstream analyses. Before downstream analysis, we applied a quality value threshold of 20 (i.e. *p*-value *≤* 0.01) for the event detection score, which is a Phred-like quality score defined as *Q* = *−*10 log_10_(*pval*). For detected events, a second quality score is available for modification identification (m6A or other modification type). Since the identification quality score depends on the methylated fraction (lower fractions yielding lower scores), we accepted identifications as m6A for a given position in all strains if the identification quality score at that position was *≥* 20 in at least one strain. We did this to avoid being over-conservative and underestimating the methylated fractions for positions which were likely to be truly methylated but with a low methylation fraction in some strains. We checked that the ability to estimate low methylation fractions for m6A was not impaired in strains with the lowest coverages by examining the relationship between coverage and m6A fraction in GATC motifs (Supplementary Figure S3).

### Genome annotation

We used the NBCI Prokaryotic Genome Annotation Pipeline (PGAP, Tatusova et al. (2016), version 2020-09-24.build4894) to annotate the assembled genome of our reference strain. The location of the origin of replication was determined using the DoriC server (Gao et al., 2012). The CDS predicted by PGAP were assembled into operons using the Operon Mapper server (http://biocomputo. ibt.unam.mx/operon_mapper/, Taboada et al. (2018), run on 2020-12-07). Gene ontology annotations for the CDS predicted by PGAP were pooled from two approaches: (i) submitting the predicted CDS to Interproscan (Jones et al. (2014), version 5.48-83.0) and (ii) running a tblastn search of the predicted CDS against all UniProtKB entries related to *Serratia marcescens* (UniProtKB database searched via https://www.uniprot.org/ on 2019-01-31) and keeping the hits with the highest bit scores.

### Identification of putative methyltransferase genes

To identify putative MTase genes in our reference strain, we downloaded the nucleotide sequences of all the entries available for the *Serratia* genus in the REBASE database (Roberts et al., 2015) (http://rebase.neb.com/rebase/rebase.html accessed on 2020-03-07) and performed a local nucleotide-to-nucleotide blast search of those sequences against the assembled genome of our reference strain. In order to identify putative MTases missing from the REBASE database, we also downloaded all the protein entries from the NCBI Identical Protein Groups (IPG) database using the search terms “DNA adenine methyltransferase” and restricting the search to the “bacteria” taxon (https://www.ncbi.nlm. nih.gov/ipg/ accessed on 2020-03-11). Those protein sequences were then matched against our reference strain genome using a local protein-to-translated nucleotide blast search. The IPG database was accessed programmatically using the rentrez R package (Winter, 2017).

### Calculation of tetramer composition bias along the genome

In order to test if the main target sequence identified for adenine methylation (5*^’^*-GATC-3*^’^*) was under differential selection depending on the genomic context, we calculated the tetramer composition bias in two sets of chromosome segments: coding sequences (CDS) and promoter regions (defined as 200-bp-long regions immediately upstream of an operon-leading CDS). Sequences for CDS were taken from their (+) strand, and sequences for promoters were taken from the strand corresponding to the (+) strand of the downstream CDS. For each of the 256 possible tetramers, we counted the total number of occurrences observed in those sets, *N*_tet_, separately for CDS and for promoter sets. We compared those observed values with the expected number of occurrences based on two models: a zero-order model based on the frequencies of each nucleotide (A, T, G, C) in each segment and assuming that tetramers were generated from a random combination of those, and a Markov chain model taking into account the underlying biases that might exist in the frequencies of dimers and trimers comprising each tetramer in a given segment. In the zero-order model, the expected number of occurrences of a given tetramer in a sequence segment *i* is (Pride et al., 2003):

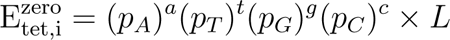

where *p_A_*, *p_T_*, *p_G_* and *p_C_* are the observed frequencies of each nucleotide in the segment, *a*, *t*, *g* and *c* are the count of each nucleotide in the considered tetramer (between 0 and 4), and *L* is the segment length. In the Markov chain approach, the expected number of occurrences of a given tetramer of composition *b*_1_*b*_2_*b*_3_*b*_4_ in a sequence segment *i* is (Rocha et al., 1998; Pride et al., 2003):

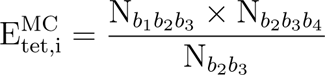

where N*_b_*_1_*_b_*_2_*_b_*_3_, N*_b_*_2_*_b_*_3_*_b_*_4_ and N*_b_*_2_*_b_*_3_ are the numbers of occurrences of the trimers *b*_1_*b*_2_*b*_3_, *b*_2_*b*_3_*b*_4_ and of the dimer *b*_2_*b*_3_ in segment *i*, respectively. For the CDS and promoter sets separately, we calculated the expected total number of occurrences as the sum of expected numbers for each segment 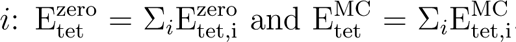. Deviations in the usage of each tetramer from expectation were then calculated as:

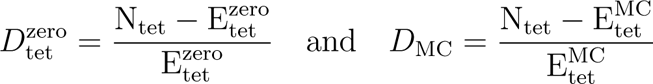

for the zero-order and the Markov chain models, respectively.

### Identification of partially methylated m6A loci in GATC motifs

GATC motifs targetted by Dam are usually methylated on both strands, but hemimethylated or non-methylated locations can be inheritable features important for the regulation of various cellular processes (Braaten et al., 1994; Casadesús and Low, 2006). To identify GATC loci which were not fully methylated in our dataset, we considered the estimated fractions of methylated adenines at each GATC locus separately for the plus and the minus strands. In any given strain, loci for which no methylated fraction was reported by the PacBio pipeline (event detection score *<* 20) were assigned a value of 0 for that strain (i.e. non-methylated). Additionally, loci for which the modification identity score was *<* 20 in all strains were also considered not to be methylated as m6A. Assuming that the distribution of modified fractions on the plus and minus strands for fully methylated loci would follow a truncated bivariate distribution centered around (1, 1) (i.e. both strands fully methylated), we defined the set of partially methylated GATC loci of interest for downstream analyses as the loci which presented modified fractions on the plus or on the minus strand which deviated from the point of full methylation at coordinates (1, 1) more than four times the average observed quadratic distance to (1, 1), in at least one strain (Supplementary Figure S4). We checked that low estimated values of methylated fractions were not a sequencing artifact due to low coverage of the corresponding GATC loci by examining the relationship between coverage bins and estimated methylated fraction (Supplementary Figure S3). We tested for association between genetic variants and methylated fractions of partially methylated m6A epiloci using Wilcoxon rank-sum tests. For each genetic variant present in more than two sequenced genomes, we ran Wilcoxon tests against all partially methylated m6A epiloci and applied a Benjamini-Hochberg false discovery rate (FDR) correction (Benjamini and Hochberg, 1995) on those tests p-values.

### Effect of evolutionary treatments on genetic and epigenetic changes

To test for an association between evolutionary treatments and genetic variants, we ran Fisher’s exact tests on the contingency tables relating genotypes and evolutionary treatments, for each genetic variant or linked variants (haplotypes) and for pooled variants (as defined in Figure 3a), with a Benjamini-Hochberg FDR correction to the tests p-values.

To determine if some adenine methylation changes had been selected by the evolutionary treatments and were stable enough to persist after several generations in common garden conditions, we tested for association between evolutionary treatments and methylated fractions of partially methylated m6A with a methylated fraction range over the sequenced clones *>* 0.2 using Kruskal-Wallis rank sum tests with Benjamini-Hochberg FDR correction. In addition to those individual tests, and to investigate whether the genes related to m6A epiloci which tended to be associated with evolutionary treatments were consistently related to some biological processes, we performed a gene ontology enrichment analysis. Each partially methylated m6A epiloci was assigned a gene if it was contained in its CDS or, alternatively, if it was located less than 500bp upstream of it. A gene assigned to an m6A epiloci was assigned this epiloci’s Kruskal-Wallis test’s uncorrected p-value. Genes assigned to more than one m6A epiloci were assigned their lowest p-values. The GO enrichment analysis was performed using the topGO R package (Alexa and Rahnenfuhrer, 2020). The reference list of genes comprised all the genes assigned a Kruskal-Wallis test’s p-value as described above, and the subset of interest was the genes which had an uncorrected p-value *<* 0.05. GO enrichment test was performed using Fisher’s exact test and the “weight01” algorithm (Alexa et al., 2006; Alexa and Rahnenfuhrer, 2020).

### Association between phenotypes and genetic and epigenetic data

We explored the association between genetic variants present in at least two sequenced genomes and phenotypic traits using Wilcoxon rank sum tests with FDR correction. We investigated the association between methylated fraction of the partially methylated m6A epiloci and phenotypic traits using Spearman’s *ρ*. We analyzed only partially methylated epiloci with a methylated fraction range *>* 0.2 and we assigned each epiloci its overlapping gene or, alternatively, the closest downstream gene within 500bp (if any).

## Results

### Genome sequences and genetic variants

We sequenced ten strains from the 31 *^◦^*C treatment, eight from the 38 *^◦^*C treatment and ten from the 24–38*^◦^*C fluctuating treatment, as well as the original reference stock strain used in the experimental evolution (Supplementary Figure S1). The reference strain chromosome was 5 117 300 bp long, with a GC content of 59.8%. The genome annotation by NCBI’s PGAP predicted 4795 CDS, organized in 2686 operons according to Operon Mapper. 52 mutations were identified from the aligned genomes of the reference and evolved strains, of which 35 were located inside coding regions and 32 were non-synonymous (Table 1 and Supplementary Figure S5). Some mutations always occurred together in the sequenced strains: we use the term “haplotype” to refer to a set of linked mutations in our dataset (Table 1). Almost all identified haplotypes were strain-specific mutations. However, a striking feature of the variant map is the presence of 11 genetic mutations associated into a single haplotype and for which the minor allele is observed in 5 of the evolved strains from 38 *^◦^*C and in the reference genome, but in no other evolved strain (haplotype *a* in Table 1). This suggests that the ancestor colony used to initiate the replicated populations in the evolution experiment, and which was itself derived from the reference strain sequenced here, actually contained at least two lineages which were preserved in some populations at 38 *^◦^*C but in no other evolutionary treatments. Even excluding variants from haplotype *a*, multiple parallel substitutions were observed among the evolved strains (Figure S5). Some genes exhibited several independent mutations in different strains: two independent mutations occurred in a deacetylase, three in a galactokinase and four in a glycosyltransferase (Table 1).

**Table 1:**
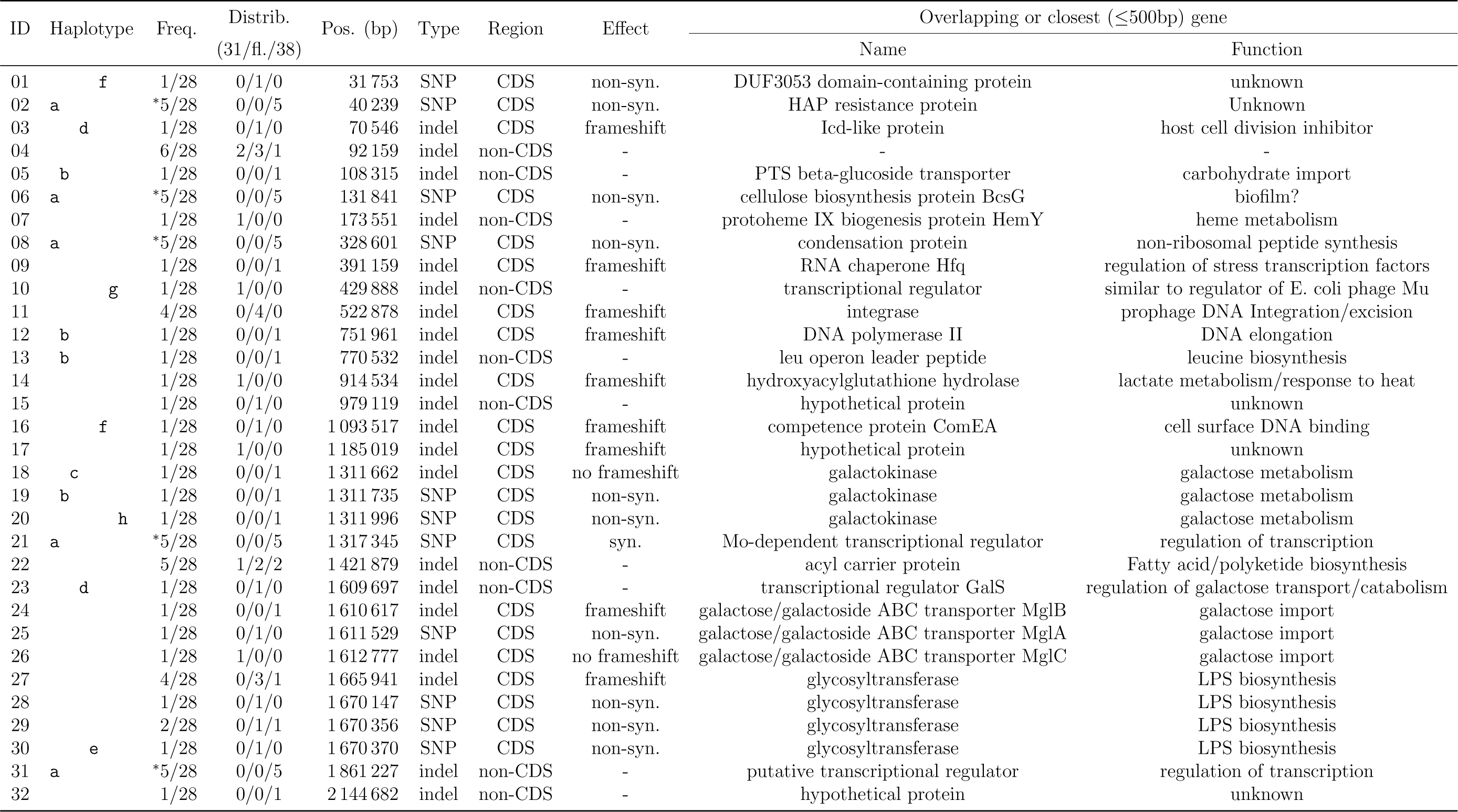
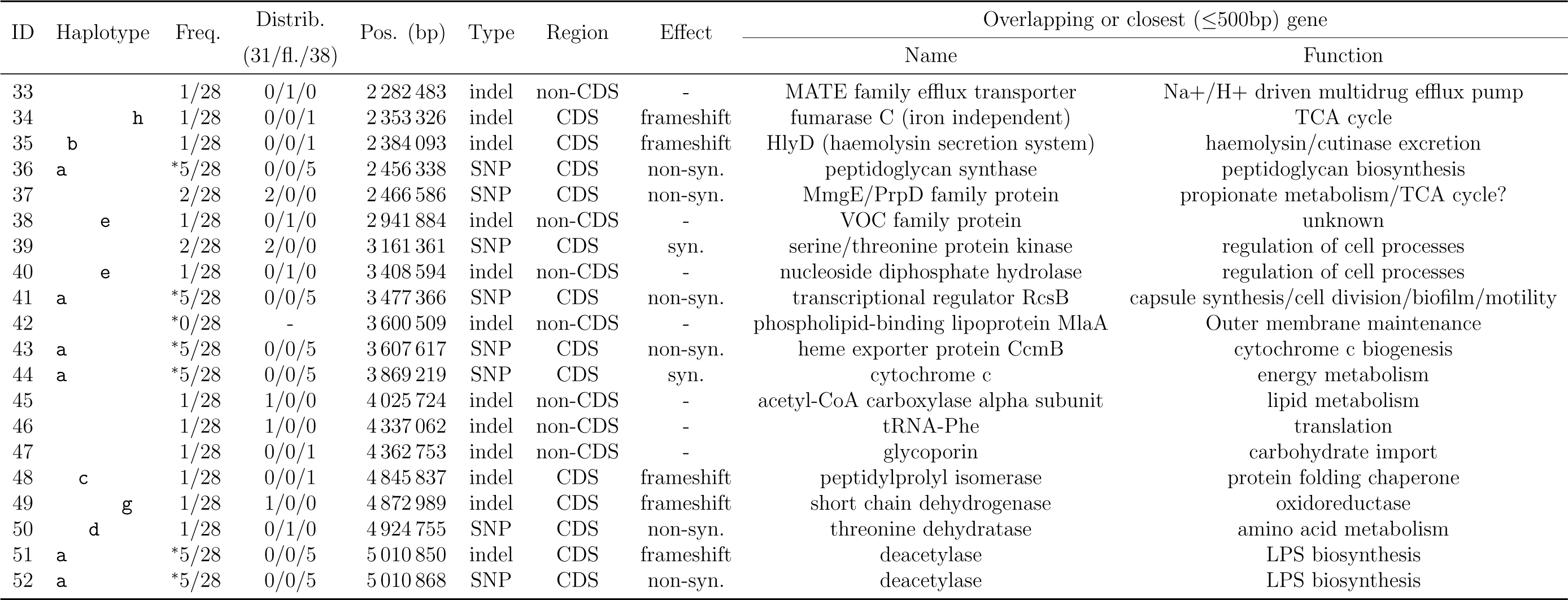
Summary of the genetic variants observed in the sequenced strains. Haplotype: letters denote groups of mutations for which alleles are associated together. Freq.: minor allele frequency among the 28 evolved strains. An asterisk denotes loci for which the reference strain carries the minor allele. Distribution of mutations per strains is shown in Supplementary Figure S5.

### Adenine methylation landscape in *Serratia marcescens*

#### Methylated bases and methylation motifs

Out of 2 057 542 adenine bases present in the bacterial chromosome, 81 434 (4.0 %) were detected as m6A (identification quality score *≥* 20 in at least one of the evolved strains or in the reference strain). Most of those m6A were in GATC motifs: out of the 81 434 positions detected as m6A, 76 228 (93.6 %) were in a GATC context. Since a total of 76 300 adenines belong to a GATC context in the genome, this corresponds to a very high rate of adenine methylation in GATC motifs: 99.9 % of GATC adenines are detected as m6A in at least one evolved strain or the reference strain and 99.7 % were detected in all sequenced genomes. When detected as modified, m6A bases had an average methylated fraction of 70.3 % (s.d. 28.3 %) outside GATC motifs and of 97.5 % (s.d. 5.0 %) inside GATC motifs. We did not find any evidence of a problematic coverage effect on the ability to estimate low methylated fraction for m6A (Supplementary Figure S3).

The PacBio pipeline identified two sets of motifs for adenine methylation (Table 2). One motif set for m6A was the GATC palindrome mentioned above and the other was the much rarer pair AAAGNNNNNNTCG / TTTCNNNNNNAGC. For both sets, almost all genomic occurrences (*>* 99 %) were detected as modified.

**Table 2:**
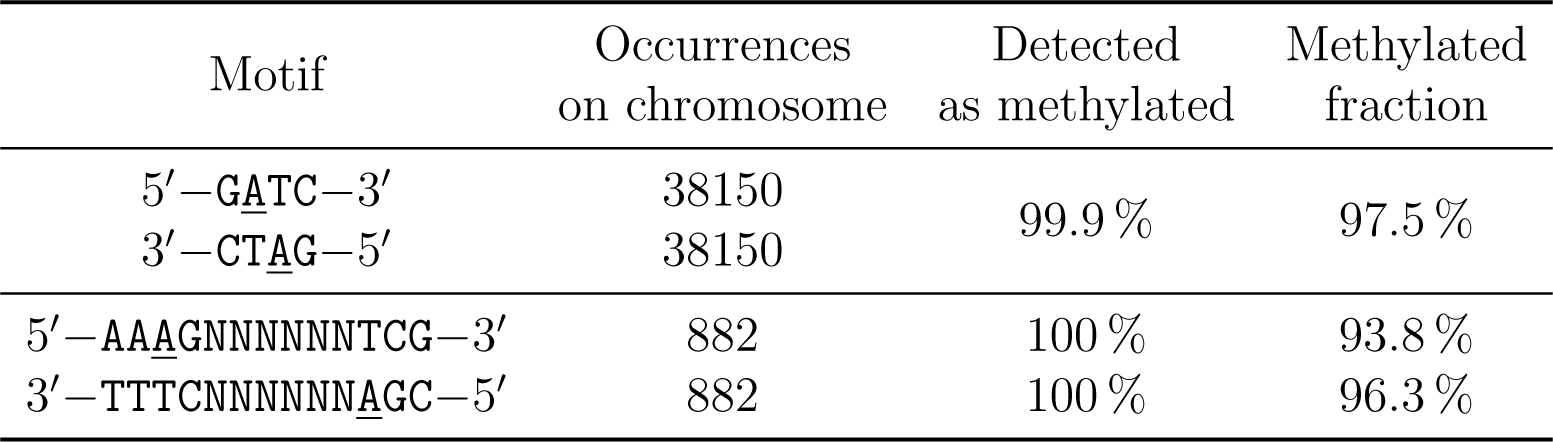
Cognate sequence motifs for m6A modification. Methylated positions are underlined. Percentages of occurrences detected as methylated are reported as mean across the 29 sequenced samples; same for methylated fractions (calculated considering only methylated positions).

Three candidate genes for adenine MTases were identified using REBASE, and no additional candidates were identified using NCBI’s IPG. One candidate was similar to M.Sma36365I, which is annotated as a type I MTase in REBASE and modifies the motif set AAGNNNNNGTTC / TTCNNNNNCAAG. It was located close to a restriction endonuclease in our reference strain genome, and is likely to be part of a restriction-modification system. A second candidate was identified as a type II MTase targetting GATC and was located inside a predicted prophage on our reference strain chromosome. Finally, the third candidate was similar to the orphan MTase M.Sma13880DamP, which also targets GATC motifs.

#### Genomic methylation profiles

We investigated m6A methylation profiles around the boundary between the first CDS of each predicted operon and the immediate upstream regions, which are likely to contain regulatory and promoter regions (Figure 1). From a base-composition perspective, upstream regions have lower GC content that coding regions, which can be related to constraints on DNA curvature or double helix stability in promoter regions (Bohlin et al., 2008) (Figure 1a). Average methylation of adenines into m6A bases was clearly lowered in upstream regions, in particular within 200 bp of the translation start sites, and reached background level again after a few tens of bp once in the operon coding regions (Figure 1b). This mostly mirrored the pattern observed for the proportion of adenines located in GATC motifs (Figure 1c).

**Figure 1:**
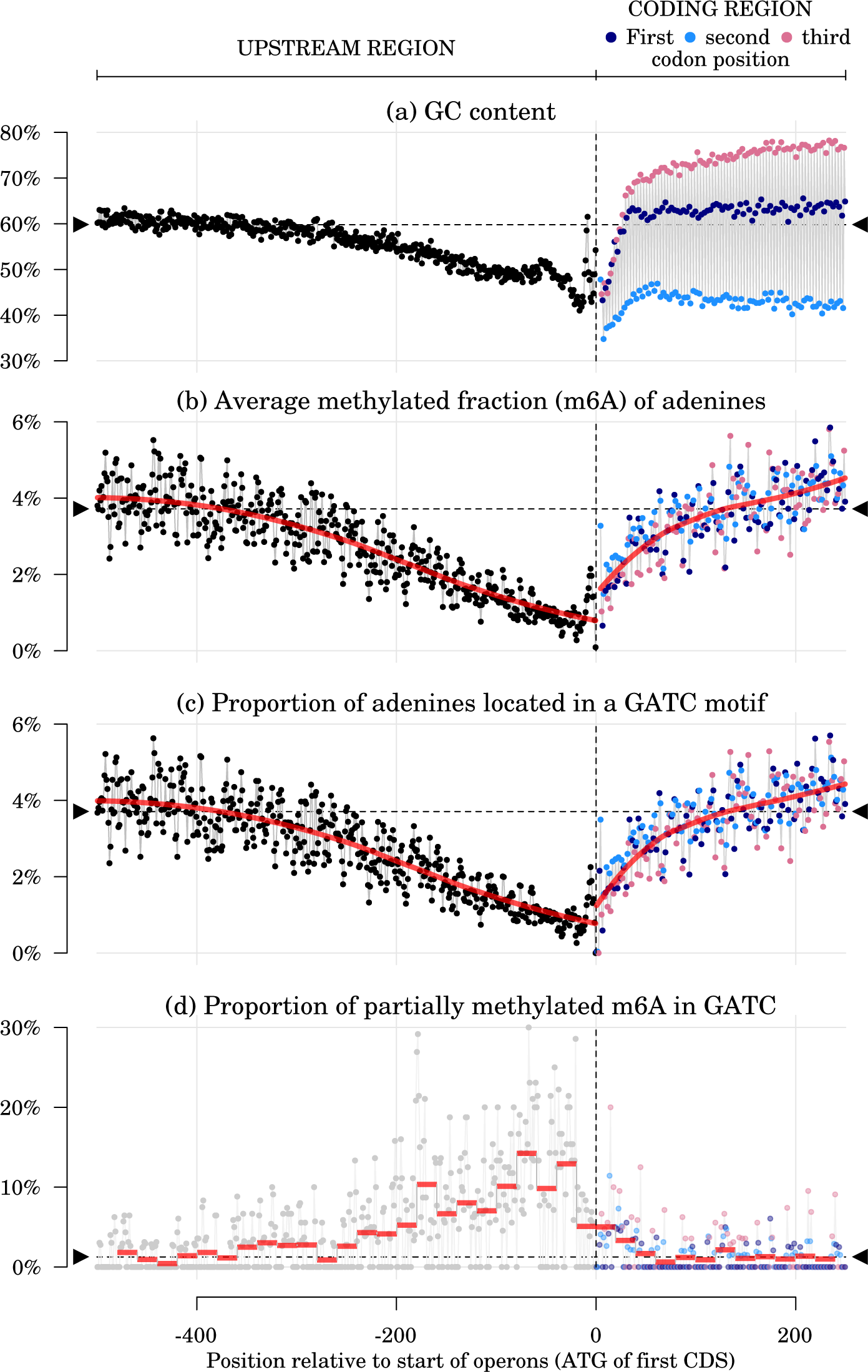
Profiles of nucleotide composition and adenine methylation around the start positions of operon-leading CDS. Red lines: LOESS regression (in (b) and (c), span = 0.75) and binned values (in (d), width = 20*bp*). Data points are averaged over each position relative to the leading CDS initiation codon, based on the 2686 operons predicted in the reference genome. Horizontal dashed lines and black triangles show genome-wide average values. Values for the three first bases on the coding sequences (usually ATG) are dropped from the plot to keep the y-scale reasonably narrow.

### GATC usage bias in *Serratia marcescens* genome

Oligonucleotide usage bias in prokaryotic genomes exists as a result of evolutionary constraints, such as codon usage bias and palindrome avoidance (Rocha et al., 1998). To investigate if any such evolutionary constraint dictated the distribution of the main *S. marcescens* adenine methyltransferase cognate motif in promoter regions, we searched for evidence of differential usage bias of the 5*^’^*-GATC-3*^’^* tetramer between coding sequences (CDS) and promoters, defined here as 200-bp long regions upstream of operon-leading CDS. In practice, we compared the usage bias of GATC with the usage bias of all other nucleotide tetramers (Figure 2). The usage bias for a tetramer in a given genomic region is positive if the tetramer is more abundant than expected by chance, and negative if it is rarer. Overall, the range of tetramer usage bias in CDS and promoters was larger when measured from a zero-order approach (deviation ranging from -0.97 to +1.95) than from a Markov chain approach (-0.71 to +1.12). The correlation between values obtained from the two approaches was moderate (Spearman’s *ρ* = 0.20), indicating that tetramer usage biases are strongly related to biases in dimers and trimers usage which are taken into account by the Markov chain approach but not by the zero-order approach. Most of the variation in tetramer usage bias was positively correlated between CDS and promoters (Spearman’s *ρ* = 0.58 for Markov chain estimates), but some tetramers exhibited large differences between their in-CDS and in-promoter usage biases (Figure 2). Remarkably, 5*^’^*-GATC-3*^’^* showed one of the largest distortions in usage bias between CDS and promoter regions among all tetramers, in both approaches. The 5*^’^*-GATC-3*^’^* tetramer was rarer in promoters and more frequent in CDS than expected by chance, even when taking into account underlying biases in dimer and trimer usage (Figure 2).

**Figure 2:**
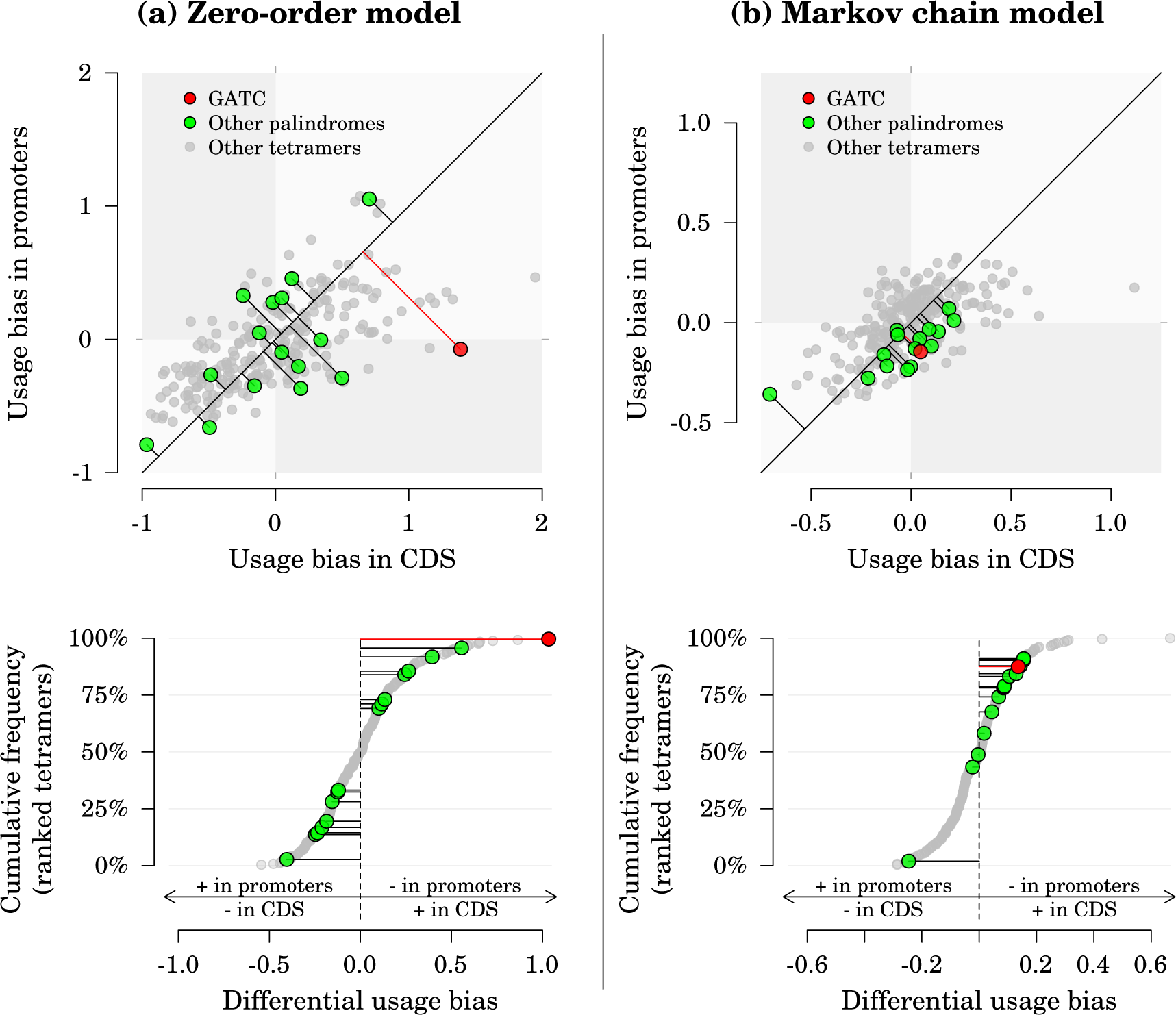
Tetramer usage bias in coding sequences (CDS) and promoters. (a) bias calculation based on observed single-nucleotide frequencies (zero-order approach). (b) bias calculation taking into account the observed frequencies of dimers and trimers (Markov chain approach). In the upper panels, the distance between a data point and its projection on the identity line measures how imbalanced the usage biases between CDS and promoters are (i.e. the differential usage bias). In the lower panels, those differential usage biases are sorted across all tetramers.

### Identification of partially methylated m6A epiloci

Non-methylated and hemimethylated GATC loci are usually rare in the genome of bacteria species possessing an adenine MTase targetting this motif. Such GATC loci can be involved in the regulation of DNA replication and of DNA-protein interactions and their methylation status can potentially be transmitted to the next generation (Braaten et al., 1994; Casadesús and Low, 2006). Here, most adenines in GATC motifs were observed close to full methylation but we identified 892 partially methylated adenines in GATC motifs, i.e. 1.2 % of all adenines in GATC motifs. They were located in 451 distinct GATC palindromes. We considered those partially methylated m6A as the m6A epiloci of interest and used them for downstream analyses of association with genetic and phenotypic data. Taking into account the genomic distribution of all GATC motifs, partially methylated m6A were more frequent than expected in promoter regions (*χ*^2^ = 1417.3, *df* = 1, *p <* 0.001) and rarer than expected in operons (*χ*^2^ = 2908.5, *df* = 1, *p <* 0.001), as is visible in Figure 1d. Finally, very little evidence of association between genetics and epigenetics was found in this experiment: based on FDR-corrected Wilcoxon rank-sum tests, associations with p-values *≤* 0.05 were only observed between haplotype *a* and five epiloci.

### Effect of evolutionary treatments on genetic and epigenetic changes

Before FDR correction, only haplotype *a* (*p <* 0.001), pooled variants for glycosyltransferase (*p* = 0.007) and for galactokinase (*p* = 0.017) and variant 11 (*p* = 0.024) were tentatively associated with evolutionary treatments (p-values *<*0.05). After FDR correction, only haplotype *a* (*p* = 0.018) remained significantly associated with evolutionary treatments, being only observed in evolved strains from the 38 *^◦^*C treatment and in the reference strain (Table 1 and Supplementary Figure S5).

Concerning the association between evolutionary treatments and partially methylated m6A epiloci, no association was significant at the *p <* 0.05 level after FDR correction. We examined if some gene ontology terms were significantly enriched in the genes which were the closest to the m6A epiloci which were tentatively associated with evolutionary treatments (uncorrected p-value for association *<* 0.05). Using Fisher’s exact test and the “weight01” algorithm (Alexa et al., 2006; Alexa and Rahnenfuhrer, 2020), five biological process (BP) categories were enriched in this set of genes (topGO’s *p <* 0.05): “cellular response to stress”, “lipopolysaccharide biosynthetic process”, “negative regulation of cellular process”, “L-amino acid transport” and “cellular component biogenesis”.

### Association between phenotypes and genotypes

We explored the association between genetic variants present in at least two sequenced genomes and phenotypic traits using Wilcoxon rank sum tests with FDR correction (Figure 3a). The only genetic variants associated with phenotypic traits at the level *p <* 0.05 were those comprising haplotype *a*, which were associated with growth rate and yield in the presence of a virus and yield at 24 *^◦^*C. Those trait values decreased in the presence of the minor allele (Figure 3b). Pooled variants for a galactokinase and a glycosyltransferase were tentatively associated with some traits (*p <* 0.1): for example, the minor alleles related to the glycosyltransferase were potentially associated with higher growth rate at 31 *^◦^*C, in the presence of DTT or in the presence of a virus.

### Association between phenotypes and partially methylated epiloci

We examined the association between methylated fraction of the partially methylated m6A epiloci and phenotypic traits using Spearman’s *ρ* and assigning the m6A epiloci to the closest genes (Figure 4). Forty genes were related to a partially methylated m6A epiloci tentatively associated with at least one trait, using an uncorrected p-value threshold of 0.005 for Spearman’s correlation. We did not apply FDR correction in this analysis, as the large number of tests run (10 traits *×* 892 partially methylated epiloci = 8920 tests) and the relatively small sample size of each test (28 evolved strains) resulted in all FDR-adjusted p-values to be *>* 0.25, even when the original uncorrected p-values were *<* 10*^−^*^4^. Gene ontology enrichment tests performed with topGO on a trait-by-trait basis revealed only very broad functional categories for genes associated with a given trait, such as metabolism, molecule transport, regulation of cellular processes, cell communication and response to chemical stimulus (Supplementary Table S1). Based on those functions, we performed a literature search for each of those forty genes to determine which had ever been found related to metabolism, nutrient uptake, cell motility and biofilm formation, or regulation of transcription, and found that most of those genes were related to at least one such category (Figure 4).

**Figure 3:**
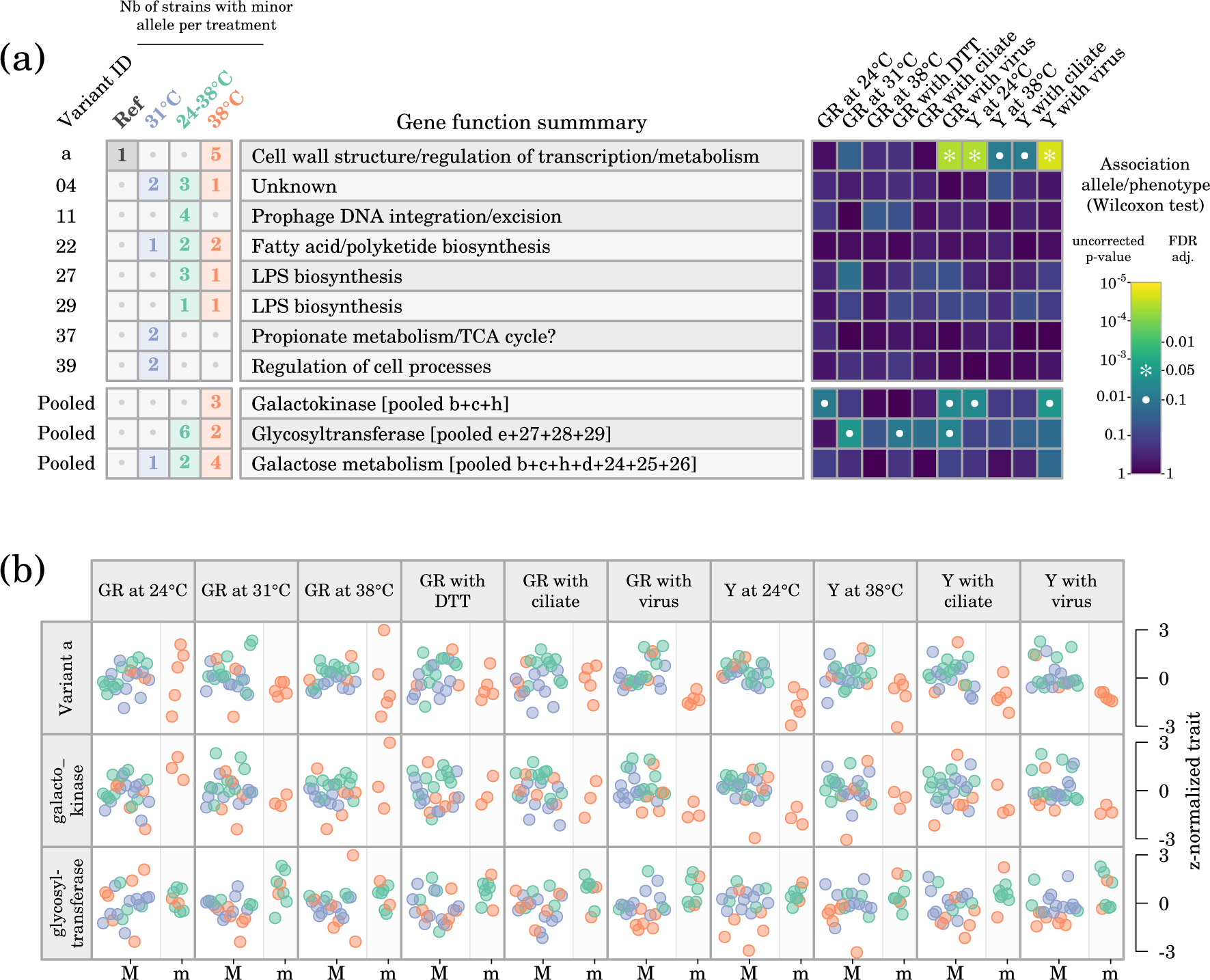
Association between genetic variants observed in at least two sequenced genomes and phenotypes. (a) Distributions of variants across evolutionary treatments and association between alleles and phenotypic traits (GR, growth rate; Y, yield). “Pooled” variants are variants affecting the same enzyme or function. (b) Detailed illustration of the association between major (M) and minor (m) alleles and phenotypes for variant *a* and for the “pooled” variants related to the galactokinase and to the glycosyltransferase. Data point colors correspond to the evolutionary treatment of each strain.

**Figure 4:**
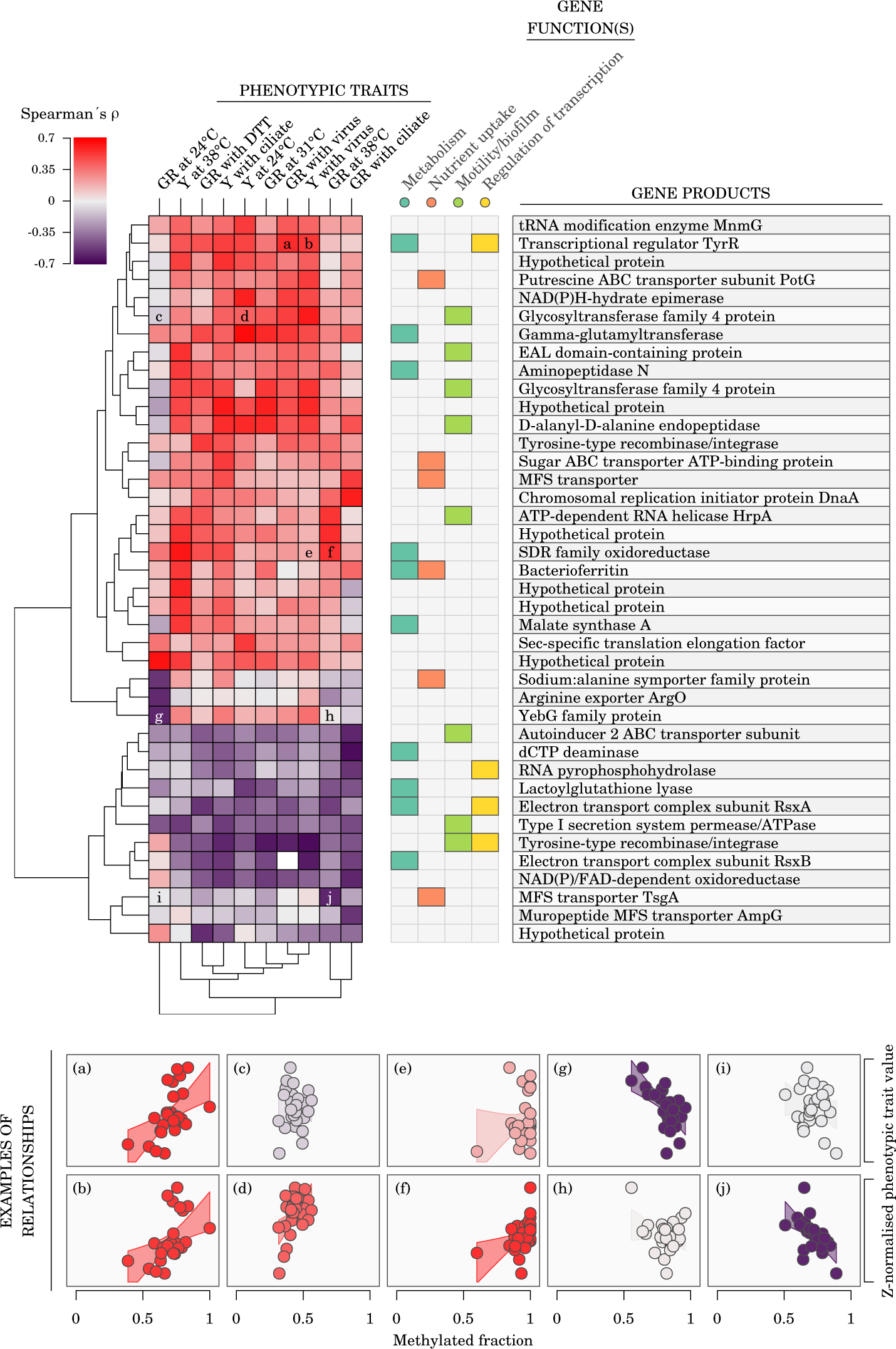
Association between adenine methylation and phenotypes. The heatmap shows Spearman’s correlation between methylated fractions of partially methylated m6A epiloci (rows) and phenotypes (columns; GR, growth rate; Y, yield) for partially methylated m6A epiloci exhibiting a methylation fraction range *≥*0.2 across sequenced samples and an uncorrected p-value *≤*0.005 for Spearman’s *ρ* with at least one trait. Overlapping or closest (*≤*500 bp) downstream genes are assigned to each epiloci. Gene functions are based on manual literature search for each gene product. Examples of relationships between m6A methylated fractions and phenotypes are given for the heatmap cells annotated with a letter.

## Discussion

We have shown that substantial variation in adenine methylation existed in *Serratia marcescens* strains after experimental evolution followed by common garden conditions, and that GATC motifs and partially methylated m6A epiloci were not randomly distributed along the genome. Sequenced strains were genetically clonal but they exhibited polymorphism in a small fraction of their m6A epiloci. In addition to de novo mutations, genetic variants were dominated by 11 linked loci (haplotype *a*) which were only observed in strains evolving at 38 *^◦^*C and suggested the existence of some degree of genetic polymorphism in the ancestor strain and differential selection between temperature treatments during the experiment. No clear effect of evolutionary treatments on specific m6A epiloci was found, suggesting that parallel environmentally-induced or environmentally-selected changes on particular epiloci were rare or labile in our experiment. Both genetic and epigenetic data were associated with some phenotypic traits to some extent. Surprisingly, the involved genes were not related to temperature response in particular but rather related to metabolism, nutrient uptake, motility/biofilm/adhesiveness and cell wall structure.

### Methylation landscape in *S. marcescens*

*E. coli* and several other Gammaproteobacteria possess a Dam enzyme responible for m6A methylation of GATC motifs (Hattman et al., 1978; Casadesús, 2016), and GATC adenine methylation in those species is involved in important functions such as protection against foreign DNA, timing of DNA replication and regulation of gene expression (Casadesús, 2016). *S. marcescens* is known to also harbour a Dam enzyme targetting GATC motifs (Ostendorf et al., 1999). The detailed *S. marcescens* adenine methylation data we collected in our experiment indeed allowed us to identify two cognate motifs in this species, including GATC. Putative adenine methyltransferases were identified in the genome, two of which could be orphan MTases responsible for GATC methylation. About 4 % of observed m6A positions were outside the identified cognate motifs, which suggests either off-target methylation by specific MTases (Murray et al., 2012), the existence of an unspecific DNA adenine MTase (Murray et al., 2018), or that some of those detected methylation events could possibly be false positives (McIntyre et al., 2019).

Dam activity in *S. marcescens* has been shown to play a role in mutation avoidance by keeping track of the reference strand during post-replication mismatch repair (Ostendorf et al., 1999). Here, several lines of evidence based on the genomic distribution of GATC motifs and of the partially methylated epiloci indicate that GATC methylation in *S. marcescens* is likely to be involved in other cell functions than solely recognition of foreign DNA or help in mismatch repair, such as gene expression regulation. Firstly, average adenine methylation was lower in regulatory and promoter regions than in the rest of the genome (Figure 1b), which was primarily due to the lower frequency of GATC motifs in those regions (Figure 1c). Secondly, this unbalanced distribution of GATC motifs was not fully accounted for by the underlying distributions of constituting di- and trinucleotides: the Markov chain model showed that GATC were rarer than expected in promoters but more frequent in CDS, suggesting that evolutionary constraints on the GATC motif differ between those genomic regions (Figure 2). Finally, partially methylated sites were not randomly distributed among the GATC sites but were more frequent in promoters and rarer in coding regions (Figure 1d). Those observations are consistent with previous results in *E. coli*, where GATC motifs are also more frequent in coding rather than non-coding regions (Barras and Marinus, 1988) and GATC located in 5*^’^* upstream regions of CDS are more frequently under-methylated (Tavazoie and Church, 1998). In *E. coli*, GATC methylation in gene regulatory regions is responsible for phase variation (Blyn et al., 1990) (but not all methylation in regulatory regions lead to regulation of gene expression -see for example van der Woude et al. (1998)). Oshima et al. (2002) suggested that GATC in upstream regions could modulate gene expression by interacting with some regulatory proteins, while Riva et al. (2004) argued that the regulation of expression was due to clusters of GATC situated inside the coding regions and which would affect DNA stability and thus expression based on their methylation status.

Finally, while each sequenced strain in this experiment appeared genetically clonal, it is important to note that the methylated fractions observed within a given strain show that most of the partially methylated m6A epiloci were heterogeneously methylated in the culture: about 55 % of the methylated fraction values for those epiloci were between 0.2 and 0.8, showing that cells in a single culture are differently methylated at those epiloci. Methylated fraction must thus be considered a culture-level property, rather than a cell-level characteristic, in the same way that bacterial phenotypic traits such as growth rate and yield are measured for a culture, not for an individual cell.

### Effect of evolutionary treatments on adenine methylation

The origin of epigenetic variation and its stability are two fundamental evolutionary questions. Changes in methylation at a genomic locus can be either genetically determined, spontaneous, or environmentally induced. In eukaryotic systems, for example, most methylation changes observed in natural plant populations generally seem to be under genetic control (Dubin et al., 2015; Hagmann et al., 2015) but only a small proportion of methylation changes were under genetic control during experimental evolution of *Chlamydomonas* algae (Kronholm et al., 2017). In eukaryotes again, both spontaneous epigenetic changes (van der Graaf et al., 2015) or environmentally induced changes (Jiang et al., 2014; Wibowo et al., 2016) have also been observed. In prokaryotes, even though the mechanisms involved in methylation processes are markedly different from the eukaryotic ones, those three types of methylation changes are also possible: genetically determined changes (e.g. mutations affecting the MTase enzyme itself or mutations in MTase target sites or in their flanking regions; Coffin and Reich (2008)), spontaneous (stochastic competition between MTase and DNA binding protein heritable via self-regulating loops), and environmentally induced (when other DNA binding proteins influencing MTase access to DNA are responding to environment, like in the case of the *pap* operon or the *agn43* gene in *E. coli* ; Wallecha et al. (2002); Casadesús and Low (2013)). Importantly, the stability of methylation states in prokaryotes can vary from a regular resetting during DNA replication to bistable epigenetic switches underpinning phase variation. For a given epigenetic switch, OFF-to-ON and ON-to-OFF rates can differ, allowing for some relative stability after a switching event (Kaminska and van der Woude, 2010; Olivenza et al., 2019). Epigenetic changes in bacteria can thus be involved in both short-term, partially heritable phenotypic plasticity and in long-term response, possibly allowing for later genetic assimilation. Given our experimental design, our conclusions below apply only to epigenetic changes that can remain stable for several generations rather than to the more labile ones.

We did not find any strong evidence to support a genetic control of adenine methylation in our experiment, as only five m6A epiloci were found associated with haplotype While the statistical power to detect association from our data is limited since there is no segregation among the bacterial clones, extensive genetic control would require assuming that each genetic mutation controls multiple different epiloci in order for genetic mutations to explain the observed epigenetic variation. Furthermore, the limited association observed between m6A epiloci and haplotype *a* could be be due to the shared history of those epiloci with haplotype *a* prior to the initiation of the evolution experiment (assuming that haplotype *a* was part of some standing genetic variation in the ancestor culture at the initiation of the experimental populations).

Any consistent methylation pattern associated with multiple clones from a given evolutionary treatment can in principle be due to methylation states induced by a common environment or explained by spontaneous changes that were environmentally selected in multiple populations, thus reflecting parallel evolution. It is also possible that some epiloci have extremely high forward- and back-mutations rates, so that some polymorphism is always present, or that the optimal fitness of a given culture is composed of a mix of subpopulations heterogeneously methylated. We did not observe any partially methylated m6A epiloci whose methylated fraction would be significantly associated with evolutionary treatments, which suggest that no parallel epigenetic changes were either induced nor selected by the evolutionary treatments, or that changes would be too labile to persist over a few generations in common garden conditions. Most of the observed epigenetic variation is thus likely to be spontaneous, stochastic fluctuations, for which our experimental design does not allow to determine if they affect the phenotype or are neutral. Highly labile epigenetic fluctuations could possibly reflect some sort of epigenetic bet-hedging mechanism (Veening et al., 2008). Our observations do not preclude, however, that environmental conditions could exert selective pressure on stochastic epigenetic variation at different epiloci that would affect a common biological process. Indeed, the results of the gene ontology enrichment analysis of the genes located by the m6A epiloci most associated with evolutionary treatments suggest that environmental pressure could act as a sieve selecting distinct epigenetic changes related to common, important biological functions.

### Distribution of genetic variants among evolutionary treatments

Most of the genetic variants observed in our experiment were present in single strains and we cannot determine if those variants provide an advantage in a specific evolutionary treatment or in the laboratory conditions in general. The only variant that was statistically associated with a particular treatment was haplotype *a*, which comprised 11 linked loci and was observed only in evolved strains from the 38 *^◦^*C treatment. Haplotype *a* was also found in the reference strain from which the ancestor used for the evolution experiment was derived. The probability of 11 mutations arising independently and in succession in several strains in our dataset is low given the duration of the evolution experiment. It is thus likely that the ancestor culture used to initiate all the populations of the experiment exhibited some genetic diversity in relation with haplotype *a*, possibly due to cell aggregation occurring during the preparation of the ancestor colony. No sign of this diversity is observed in the sequenced strains evolved at 31 *^◦^*C and 24–38*^◦^*C, indicating that haplotype *a* was driven to low frequencies or extinction in 31 *^◦^*C and 24–38*^◦^*C conditions. This indicates that evolving at lower average temperature might impose stronger selective constraints on the genetic variants in our experimental setup while higher temperature allowed for more diverse genetic trajectories.

Interestingly, we observed mutations occurring independently at different positions within several genes and being selected in parallel. This happened independently of haplotype *a* for a galactokinase and for a glycosyltransferase. Given the large effective population size in our experiment (*N_e_* = 2.6 *×* 10^6^, see Methods), genetic drift cannot explain the fixation of multiple parallel mutations in independent populations, suggesting instead that selection favoured a modified function for those particular genes. Remarkably, those two genes and the deacetylase gene (with two linked mutations belonging to haplotype *a*) are all involved in the biosynthesis of lipopolysaccharide. Mutations in those three genes were observed in the 24–38*^◦^*C and 38 *^◦^*C treatments, but not in the 31 *^◦^*C treatment.

### Effect of genetic and epigenetic variation on phenotypes

Our main objective was to determine the potential contributions of genetics and epigenetics to adaptation in rapidly changing environments. In the original evolution experiment during which the evolved strains were generated, 12 colonies were isolated from each replicate population, thus providing good statistical power to detect an effect of evolutionary treatment on phenotypic traits and enabling to show that strains from the fluctuating 24–38*^◦^*C treatment outperformed strains that evolved at constant 31 *^◦^*C (Ketola et al., 2013). In the subset of strains we sequenced here, phenotypic variability results in overlapping phenotype ranges across treatments (Supplementary Figure S2). Given this overlap, trying to link genetic and epigenetic data directly to phenotypic trait values rather than to evolutionary treatments might give insight into how genetic variants and partially methylated m6A epiloci can influence phenotype. Note that given our experimental design, where evolved clones were isolated and cultured in common garden conditions before phenotypic measurements and DNA extraction, short-lived epigenetic changes involved in rapid phenotypic plasticity would be lost and only more persistent changes, stable over several generations and more amenable to playing a role in evolution, would be carried over from the evolution experiment.

Evidence supporting a potential effect on phenotypes was found both for genetic variants and for partially methylated m6A epiloci. Given that most genetic variants were present in only one strain, the ability to detect statistically significant associations was limited, but haplotype *a* was associated with some reduced performance in the strains evolved at 38 *^◦^*C and independent genetic variants associated with galactokinase and glycosyltransferase were tentatively associated with phenotypic changes. From a general biological perspective, those genes and the genes affected by haplotype *a* are somehow related to cell wall structure, metabolism and regulation of transcription, which are all crucial cell functions likely to be impacted when environmental conditions change, but not to thermal stress in particular, and were associated both with temperature-related traits and with performances in novel environments (exposure to DTT, ciliate or virus). Similarly, partially methylated m6A epiloci which were found to be associated with phenotypes were related to genes involved in important, broad functional categories (metabolism, nutrient uptake, motility and biofilm formation). Metabolism and nutrient uptake are both likely to be crucial in determining growth rate and yield, especially after evolving in a nutrient-limiting medium such as SPL 1%.

## Conclusion

We have shown that substantial epigenetic variation existed in adenine methylation in *Serratia marcescens*, both across and within cultures of evolved strains. The distribution of methylated fractions and of the main adenine MTase cognate motif along the genome pointed to a probable connection between adenine methylation and gene expression regulation, as is the case in other Gammaproteobacteria. No clear evidence of a genetic control of epigenetic variation, nor of stable environmentally-induced or environmentally-selected epigenetic changes were found in our experiment. However, this result does not rule out the existence of labile, short-term epigenetic responses in particular in fluctuationg environmental conditions, which could be explored with more massive methylome sequencing of evolved strains grown in their treatment conditions and challenged in new conditions. Genetic variants from unexpected pre-existing standing genetic variation at the beginning of the evolution experiment seemed to be responsible for the majority of divergent phenotypic adaptation between the 38 *^◦^*C treatment and the others, but indications of both genetic and epigenetic effects on some of the phenotypic traits observed in our experiment were found.

## Acknowledgements

We acknowledge the Academy of Finland (Project 278751) and the Centre of Excellence in Biological Interactions for funding and facilities, and CSC-IT center for Science for computational resources.

## Authors contributions

TK and IK conceptualized the study. Experimental design for PacBio sequencing was done by TK and MB. DNA extraction for PacBio sequencing was done by RA. MB, IK and TK analyzed the data and wrote the original draft with later review by all co-authors.

## Data availability

PacBio sequencing data (HDF5 files) were submitted to the European Nucleotide Archive’s Sequence Read Archive (ENA-SRA) (https://www.ebi.ac. uk/ena, project PRJEB40306) and assembled genomes to NCBI’s GenBank (biosamples SAMEA7301478 to SAMEA7301506). Genetic, epigenetic and phenotypic datasets used for analysis will be submitted to the Dryad repository (https://datadryad.org).

## Supplementary Tables

**Supplementary Table S1:**
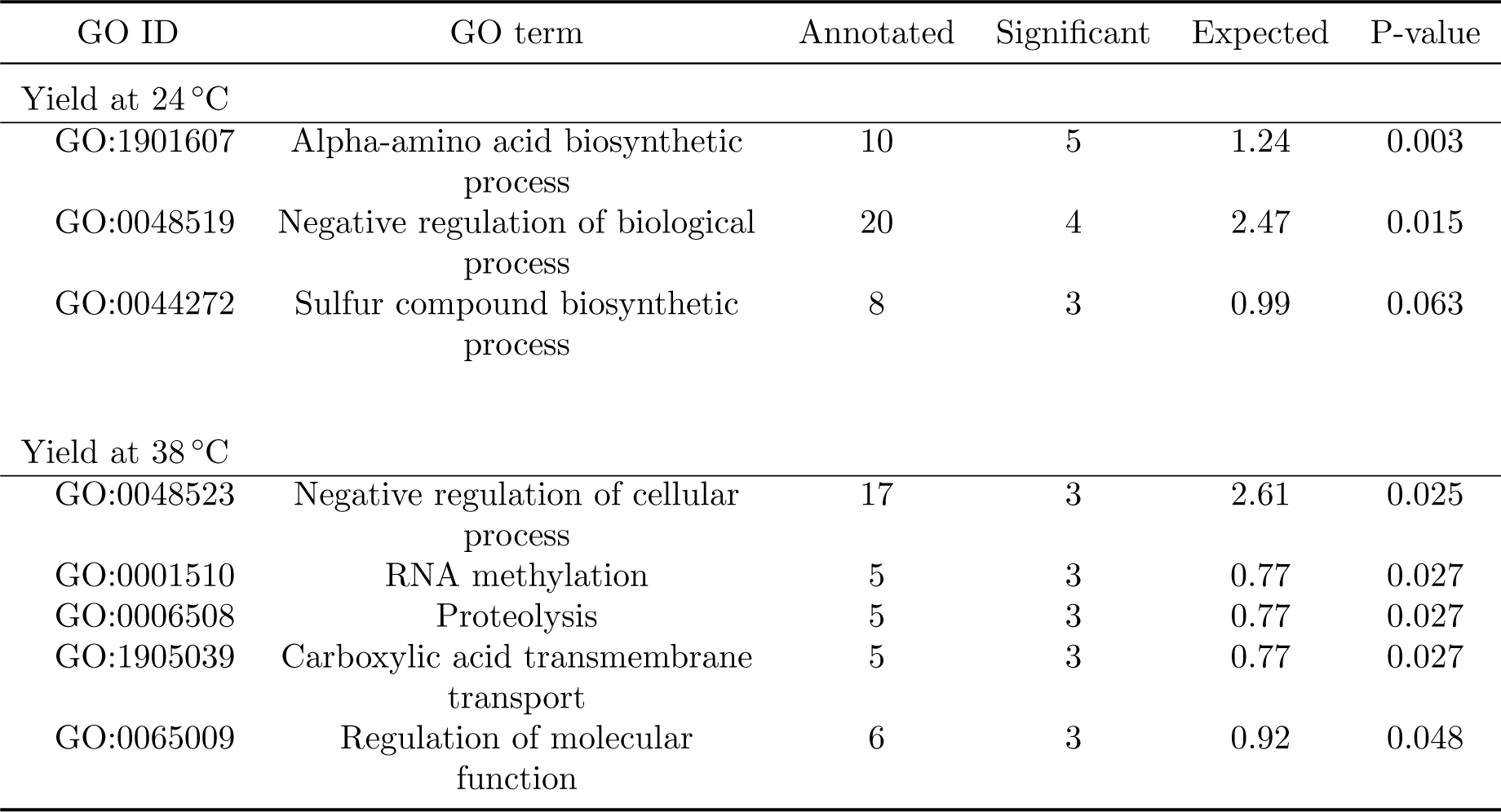
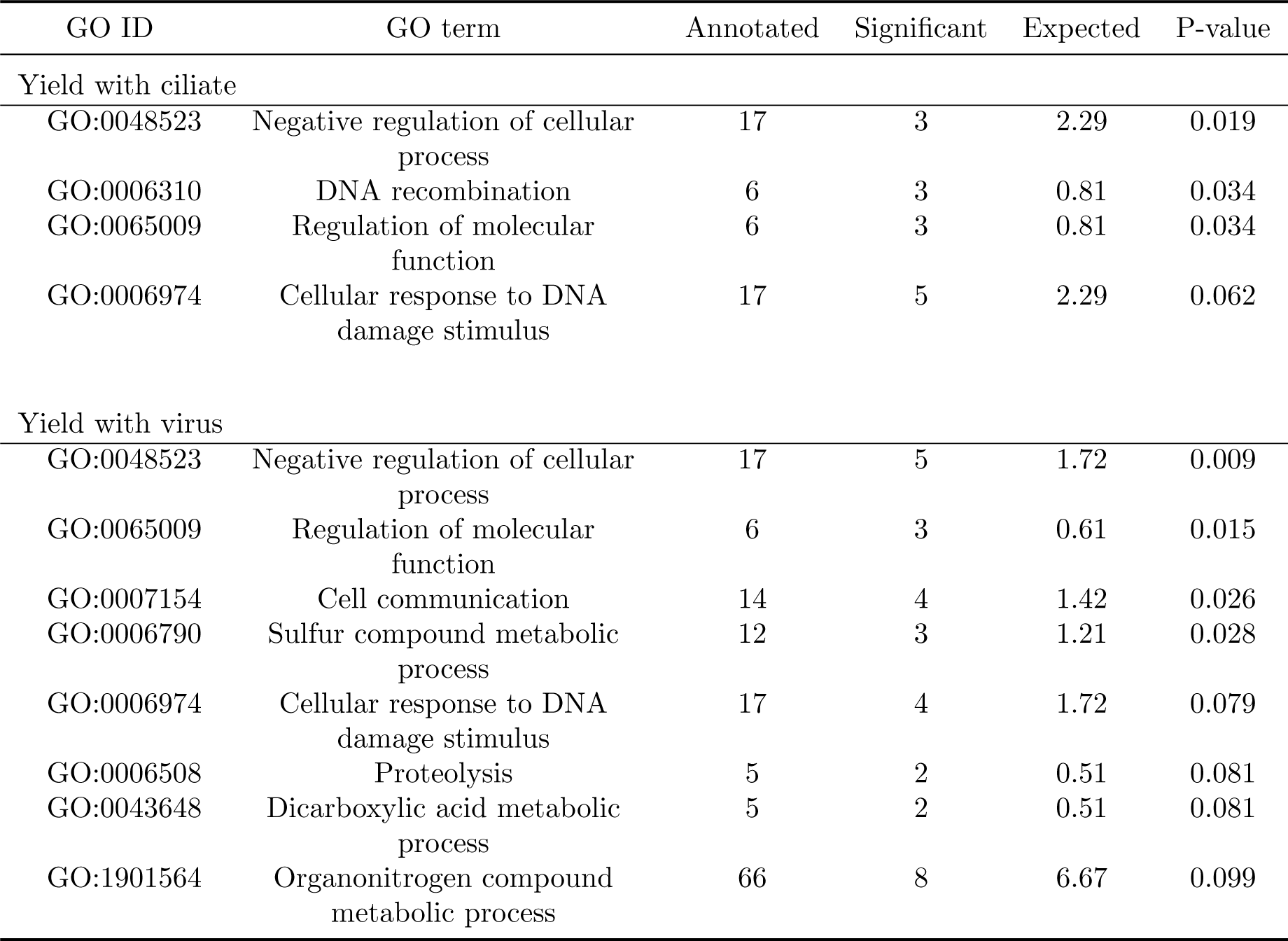
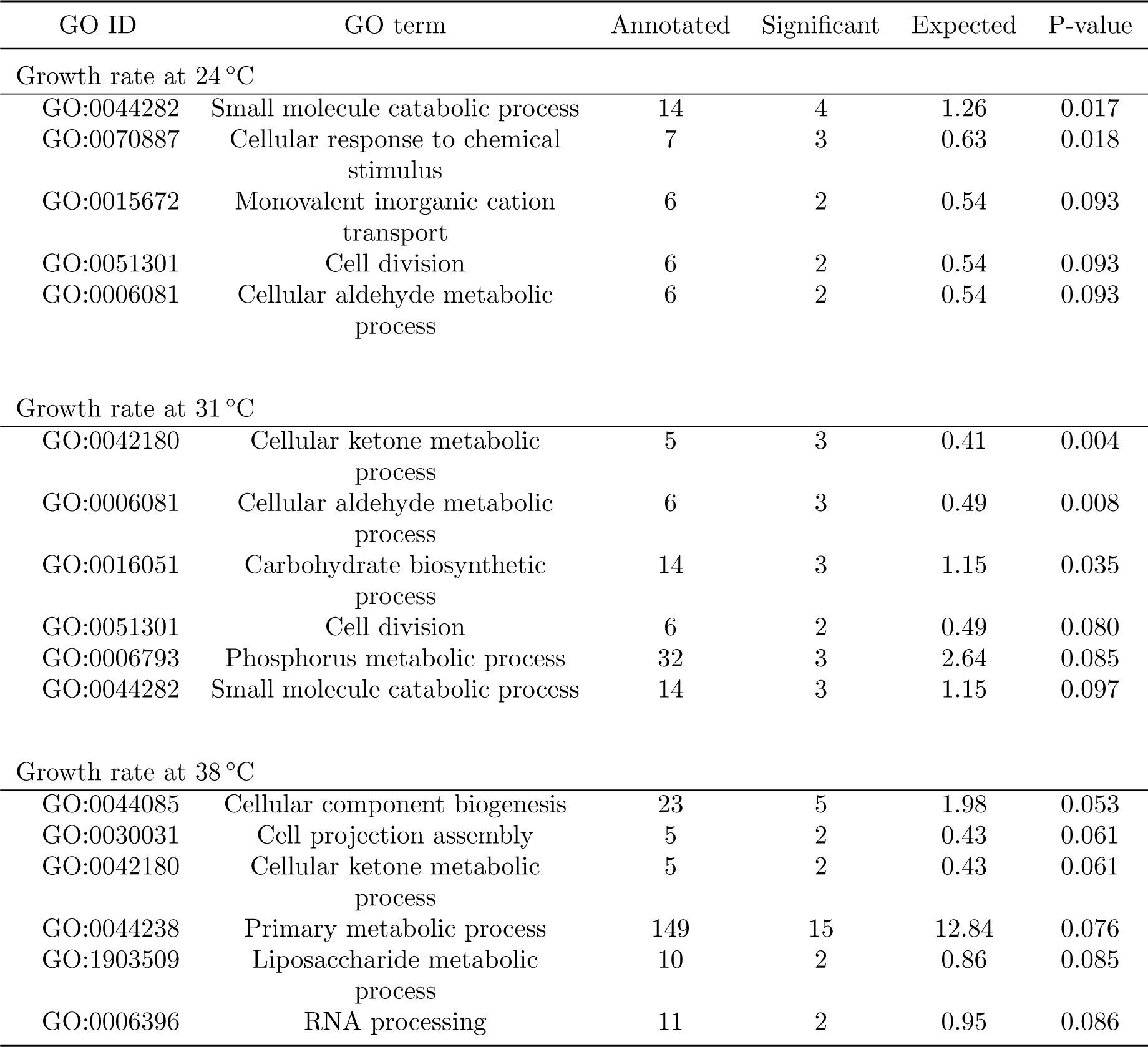
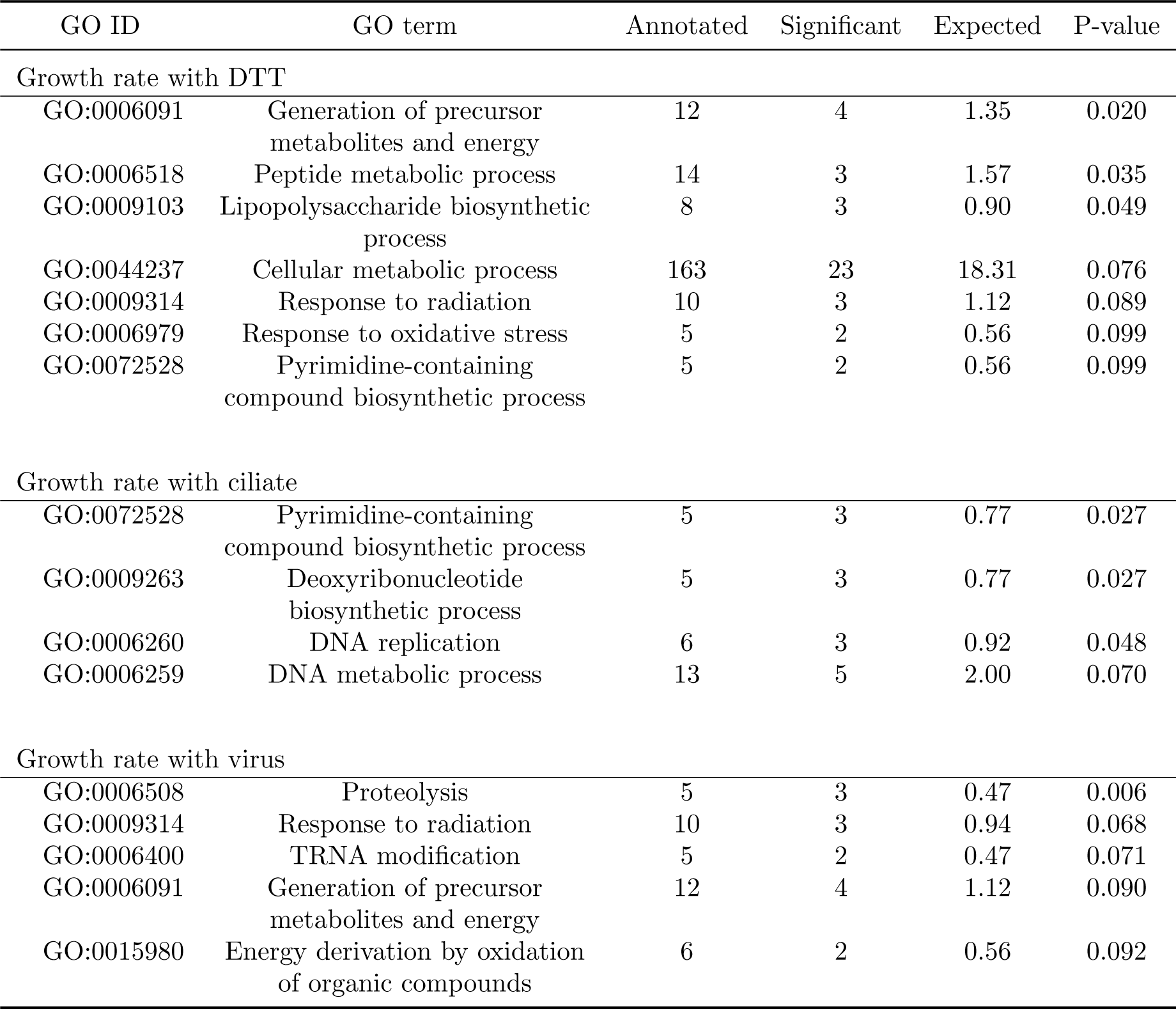
Results of the gene ontology (GO) enrichment tests based on the association between m6A methylation and phenotypes. Partially methylated m6A epiloci exhibiting a methylation fraction range ≥0.2 across sequenced samples were considered and the overlapping or closest (≤500 bp) downstream gene, if any, was assigned to each of those epiloci. For each phenotypic trait and each epiloci, Spearman’s ρ was calculated between the trait values and the epiloci methylated fractions, and the corresponding uncorrected p-value was assigned to the gene associated to the epiloci. We then defined the set of genes of interest based on a threshold of Spearman’s p = 0.05 and performed a GO enrichment test using topGO with the Fisher’s exact test and the “weight01” algorithm (Alexa et al., 2006; Alexa and Rahnenfuhrer, 2020). GO terms with enrichment test *p* ≤ 0.1 are reported for each trait (table continued on next page).

| GO ID | GO term | Annotated | Significant | Expected | P-value |
| --- | --- | --- | --- | --- | --- |
| Yield at 24 °C |  |  |  |  |  |
| GO:1901607 | Alpha-amino acid biosynthetic process | 10 | 5 | 1.24 | 0.003 |
| GO:0048519 | Negative regulation of biological process | 20 | 4 | 2.47 | 0.015 |
| GO:0044272 | Sulfur compound biosynthetic process | 8 | 3 | 0.99 | 0.063 |
| Yield at 38 °C |  |  |  |  |  |
| GO:0048523 | Negative regulation of cellular process | 17 | 3 | 2.61 | 0.025 |
| GO:0001510 | RNA methylation | 5 | 3 | 0.77 | 0.027 |
| GO:0006508 | Proteolysis | 5 | 3 | 0.77 | 0.027 |
| GO:1905039 | Carboxylic acid transmembrane transport | 5 | 3 | 0.77 | 0.027 |
| GO:0065009 | Regulation of molecular function | 6 | 3 | 0.92 | 0.048 |
| Yield with ciliate |  |  |  |  |  |
| GO:0048523 | Negative regulation of cellular process | 17 | 3 | 2.29 | 0.019 |
| GO:0006310 | DNA recombination | 6 | 3 | 0.81 | 0.034 |
| GO:0065009 | Regulation of molecular function | 6 | 3 | 0.81 | 0.034 |
| GO:0006974 | Cellular response to DNA damage stimulus | 17 | 5 | 2.29 | 0.062 |
| Yield with virus |  |  |  |  |  |
| GO:0048523 | Negative regulation of cellular process | 17 | 5 | 1.72 | 0.009 |
| GO:0065009 | Regulation of molecular function | 6 | 3 | 0.61 | 0.015 |
| GO:0007154 | Cell communication | 14 | 4 | 1.42 | 0.026 |
| GO:0006790 | Sulfur compound metabolic process | 12 | 3 | 1.21 | 0.028 |
| GO:0006974 | Cellular response to DNA damage stimulus | 17 | 4 | 1.72 | 0.079 |
| GO:0006508 | Proteolysis | 5 | 2 | 0.51 | 0.081 |
| GO:0043648 | Dicarboxylic acid metabolic process | 5 | 2 | 0.51 | 0.081 |
| GO:1901564 | Organonitrogen compound metabolic process | 66 | 8 | 6.67 | 0.099 |
| Growth rate at 24 °C |  |  |  |  |  |
| GO:0044282 | Small molecule catabolic process | 14 | 4 | 1.26 | 0.017 |
| GO:0070887 | Cellular response to chemical stimulus | 7 | 3 | 0.63 | 0.018 |
| GO:0015672 | Monovalent inorganic cation transport | 6 | 2 | 0.54 | 0.093 |
| GO:0051301 | Cell division | 6 | 2 | 0.54 | 0.093 |
| GO:0006081 | Cellular aldehyde metabolic process | 6 | 2 | 0.54 | 0.093 |
| Growth rate at 31 °C |  |  |  |  |  |
| GO:0042180 | Cellular ketone metabolic process | 5 | 3 | 0.41 | 0.004 |
| GO:0006081 | Cellular aldehyde metabolic process | 6 | 3 | 0.49 | 0.008 |
| GO:0016051 | Carbohydrate biosynthetic process | 14 | 3 | 1.15 | 0.035 |
| GO:0051301 | Cell division | 6 | 2 | 0.49 | 0.080 |
| GO:0006793 | Phosphorus metabolic process | 32 | 3 | 2.64 | 0.085 |
| GO:0044282 | Small molecule catabolic process | 14 | 3 | 1.15 | 0.097 |
| Growth rate at 38 °C |  |  |  |  |  |
| GO:0044085 | Cellular component biogenesis | 23 | 5 | 1.98 | 0.053 |
| GO:0030031 | Cell projection assembly | 5 | 2 | 0.43 | 0.061 |
| GO:0042180 | Cellular ketone metabolic process | 5 | 2 | 0.43 | 0.061 |
| GO:0044238 | Primary metabolic process | 149 | 15 | 12.84 | 0.076 |
| GO:1903509 | Liposaccharide metabolic process | 10 | 2 | 0.86 | 0.085 |
| GO:0006396 | RNA processing | 11 | 2 | 0.95 | 0.086 |
| Growth rate with DTT |  |  |  |  |  |
| GO:0006091 | Generation of precursor metabolites and energy | 12 | 4 | 1.35 | 0.020 |
| GO:0006518 | Peptide metabolic process | 14 | 3 | 1.57 | 0.035 |
| GO:0009103 | Lipopolysaccharide biosynthetic process | 8 | 3 | 0.90 | 0.049 |
| GO:0044237 | Cellular metabolic process | 163 | 23 | 18.31 | 0.076 |
| GO:0009314 | Response to radiation | 10 | 3 | 1.12 | 0.089 |
| GO:0006979 | Response to oxidative stress | 5 | 2 | 0.56 | 0.099 |
| GO:0072528 | Pyrimidine-containing compound biosynthetic process | 5 | 2 | 0.56 | 0.099 |
| Growth rate with ciliate |  |  |  |  |  |
| GO:0072528 | Pyrimidine-containing compound biosynthetic process | 5 | 3 | 0.77 | 0.027 |
| GO:0009263 | Deoxyribonucleotide biosynthetic process | 5 | 3 | 0.77 | 0.027 |
| GO:0006260 | DNA replication | 6 | 3 | 0.92 | 0.048 |
| GO:0006259 | DNA metabolic process | 13 | 5 | 2.00 | 0.070 |
| Growth rate with virus |  |  |  |  |  |
| GO:0006508 | Proteolysis | 5 | 3 | 0.47 | 0.006 |
| GO:0009314 | Response to radiation | 10 | 3 | 0.94 | 0.068 |
| GO:0006400 | TRNA modification | 5 | 2 | 0.47 | 0.071 |
| GO:0006091 | Generation of precursor metabolites and energy | 12 | 4 | 1.12 | 0.090 |
| GO:0015980 | Energy derivation by oxidation of organic compounds | 6 | 2 | 0.56 | 0.092 |

## Supplementary Figures

**Supplementary Figure S1:**
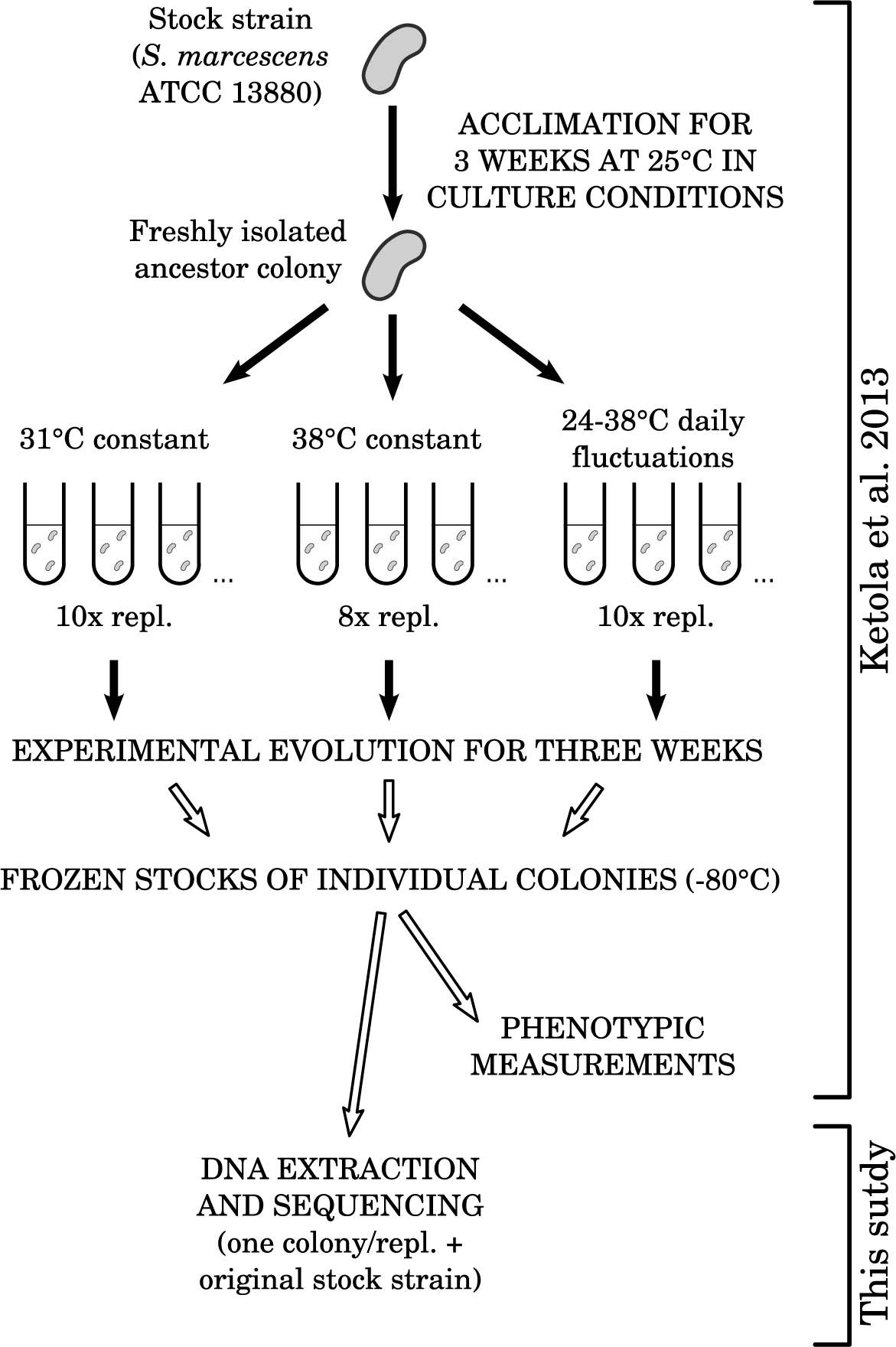
Setup of the evolution experiment from which sequenced clones were isolated. Open arrows after experimental evolution indicates steps where evolved clones were grown under common garden conditions.

**Supplementary Figure S2:**
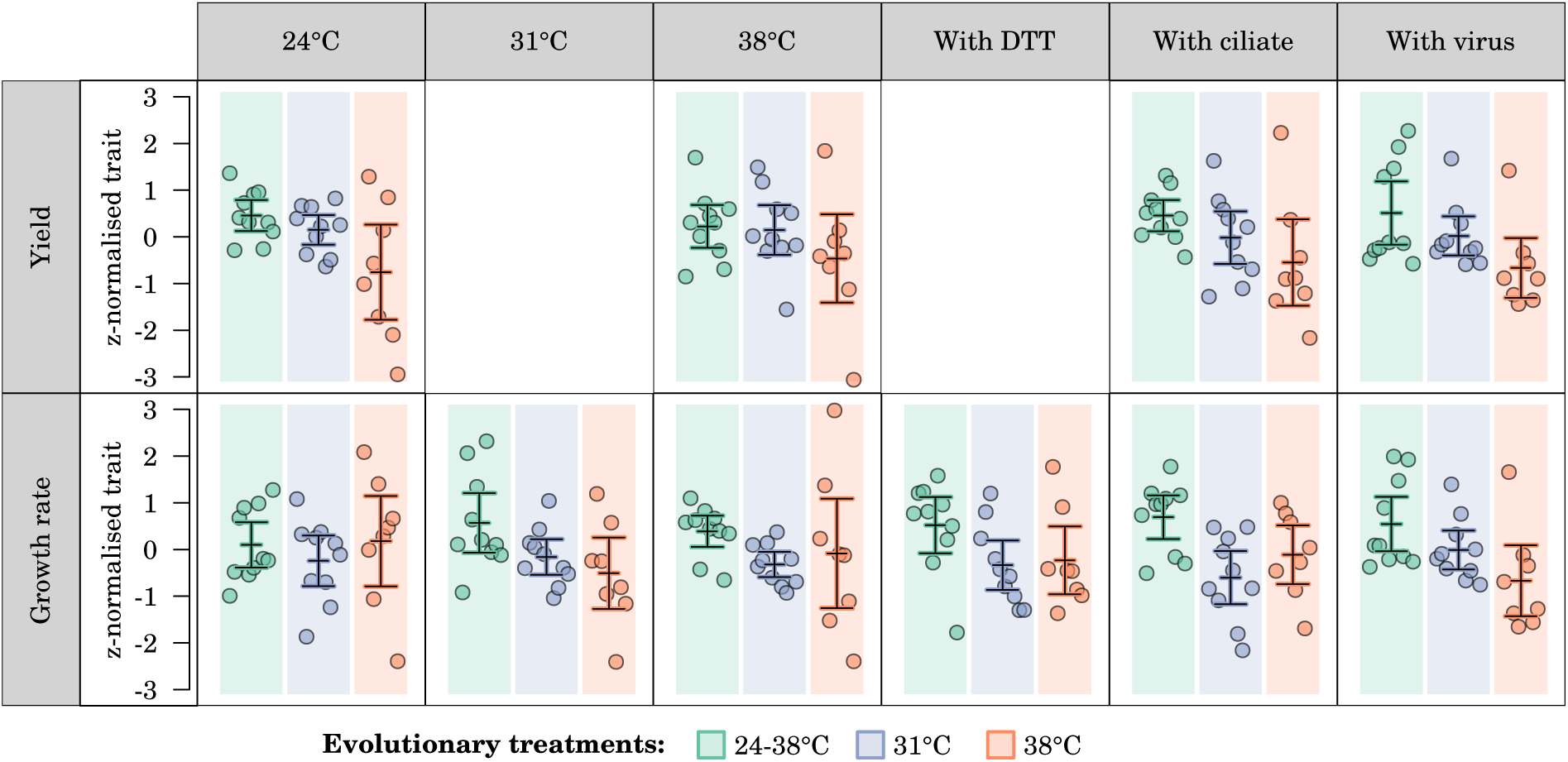
Phenotypic traits measured in Ketola et al. (2013) and used in the present study. Each data point represents one evolved strain sequenced in this study. Bars and whiskers are mean values +/- × .96 s.e. for all sequenced strains within an evolutionary treatment.

**Supplementary Figure S3:**
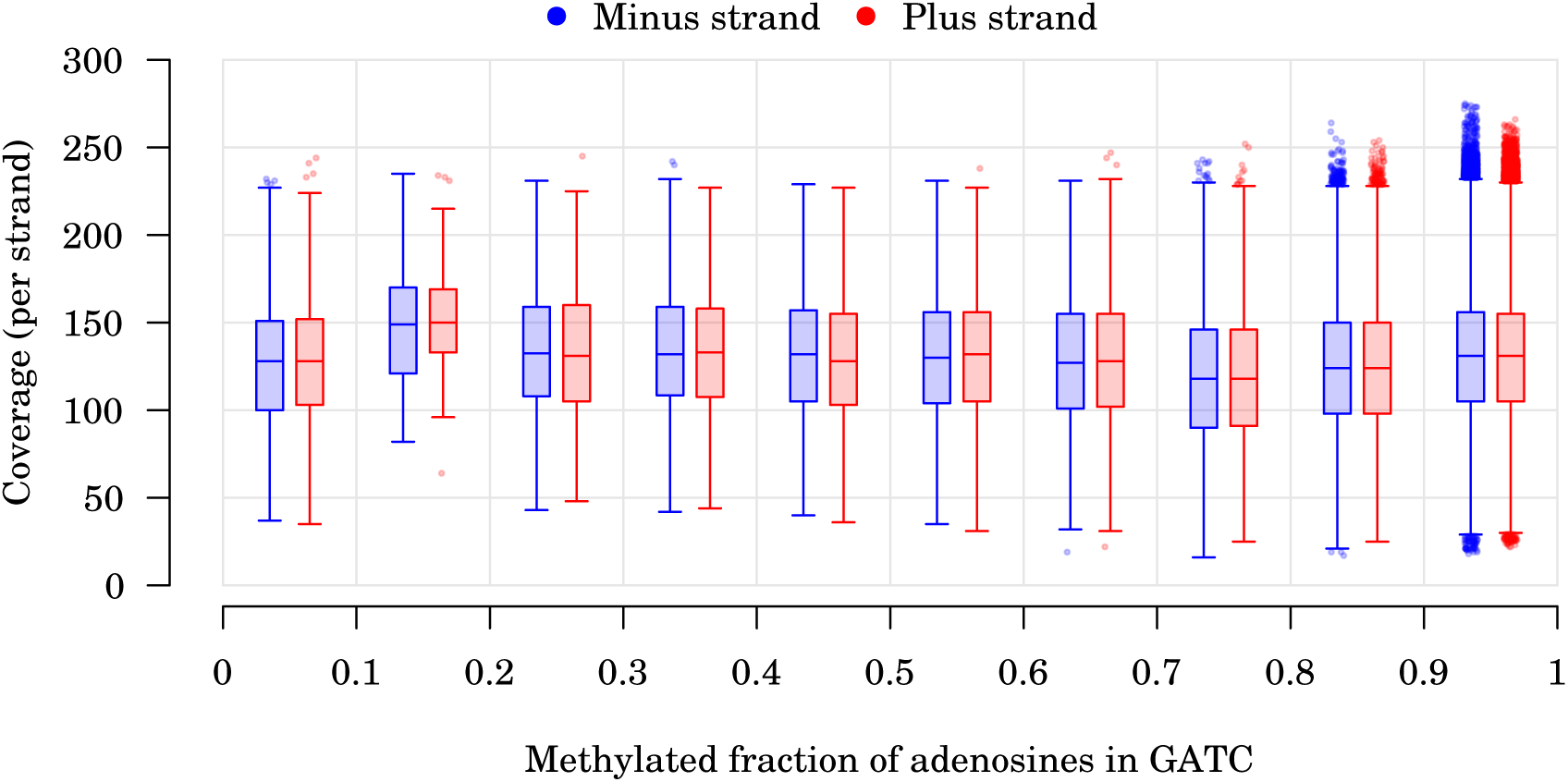
Relationship between coverage and estimated fraction of methylated adenines in GATC motifs. All adenines present in GATC motifs, from all sequenced strains, are included. Data is binned by intervals of methylated fraction of 0.1 width, as indicated on the *x* axis. There is no correlation between coverage and estimated methylated fraction (Spearman’s *ρ* = -0.01). However, the slight increase in average coverage for the (0.1, 0.2) methylated fractions compared to the (0, 0.1) fractions suggests that at very low methylation levels (below 0.2), higher coverage is needed to estimate a methylated fraction other than 0.

**Supplementary Figure S4:**
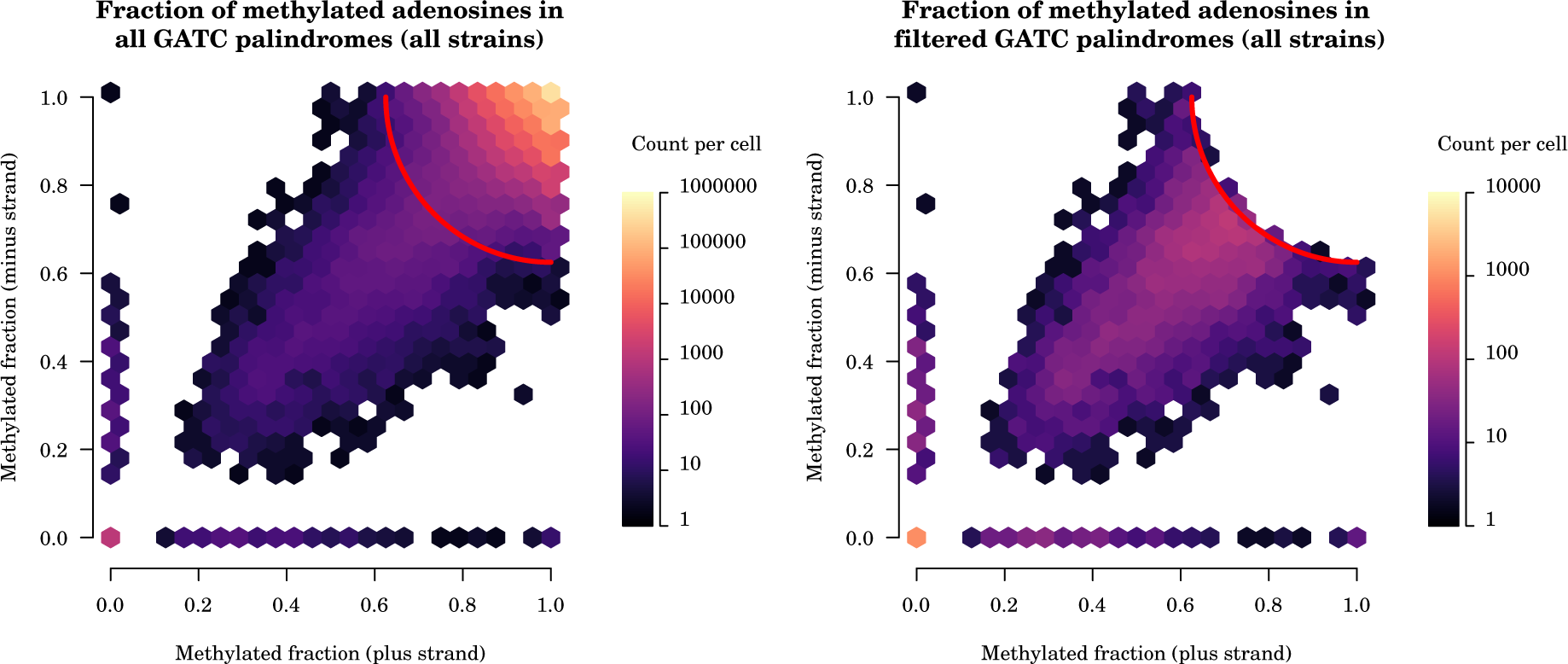
Detection of partially methylated GATC loci. Distribution of methylated fractions of adenines on boths DNA strands for GATC palindromes (showing data for all strains together). Left panel, all GATC palindromes shown; right panel, only GATC palindromes qualified as low methylation sites shown. The red arc in the left panel delimits the observations which are less four times the average quadratic distance to full methylation (point at (1,1)) away from full methylation. GATC palindromes are considered as low methylation sites if they lay outside this area (right panel).

**Supplementary Figure S5:**
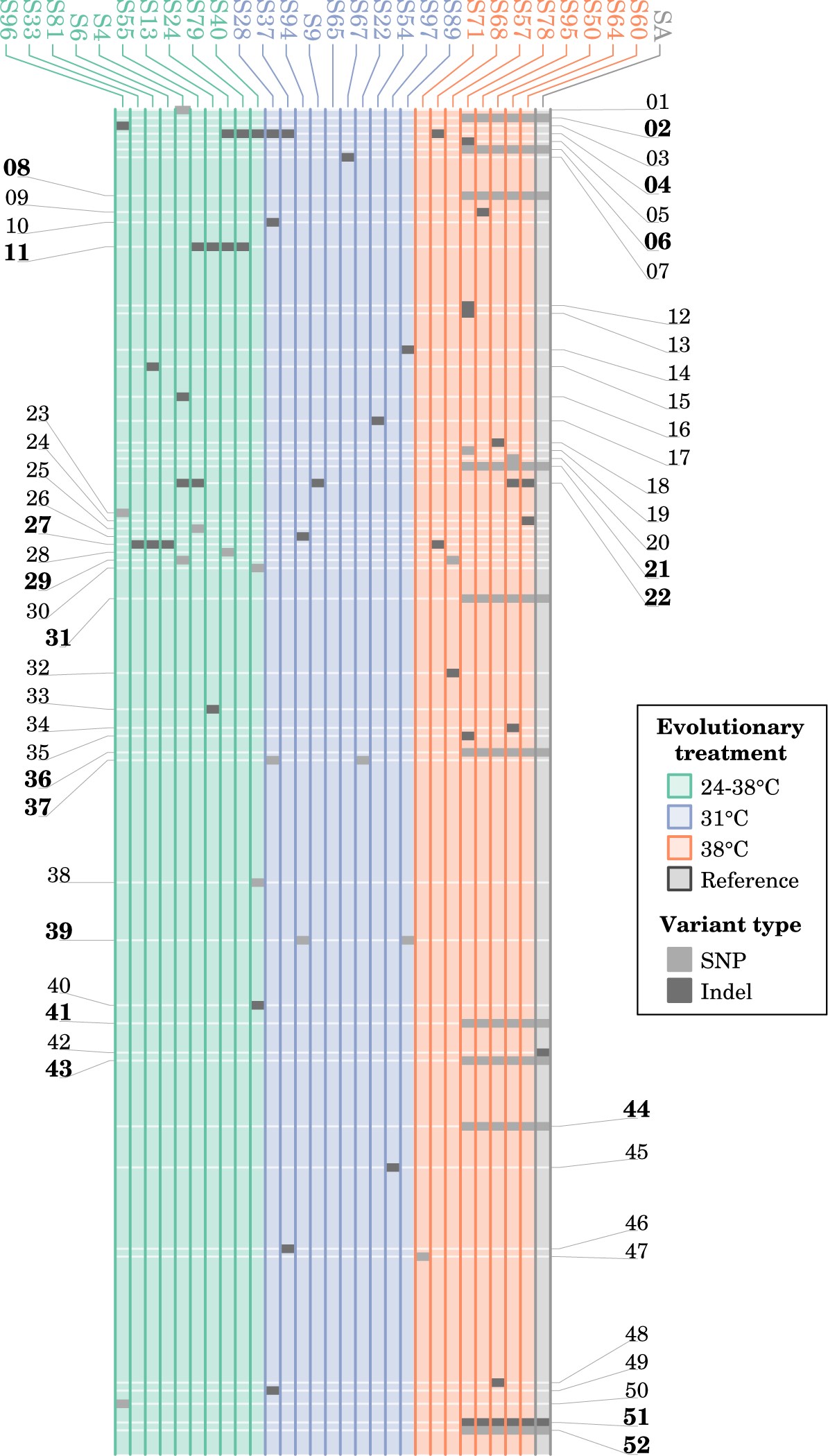
Distribution of genetic variants among sequenced *S. marcescens* clones. Each vertical lane represents the chromosome sequence of one clone. The locations along the chromosome are approximate in order to keep close loci visually distinct. Minor alleles of genetic variants are depicted. The numerical IDs of variants observed in more than one genome are shown in bold. Details about each variant are available in Table 1.

